# Bacterial lifestyle switch in response to algal metabolites

**DOI:** 10.1101/2022.01.10.475657

**Authors:** Noa Barak-Gavish, Bareket Dassa, Constanze Kuhlisch, Inbal Nussbaum, Gili Rosenberg, Roi Avraham, Assaf Vardi

**Affiliations:** Department of Plant and Environmental Sciences, Weizmann Institute of Science, Rehovot 7610001, Israel; Bioinformatics Unit, Department of Life Sciences Core Facilities, Weizmann Institute of Science, Rehovot 7610001, Israel; Department of Biological Regulation, Weizmann Institute of Science, Rehovot 7610001, Israel

**Keywords:** Chemical communication, Phytoplankton, Roseobacter, DMSP, Benzoate.

## Abstract

Unicellular algae, termed phytoplankton, greatly impact the marine environment by serving as the basis of marine food webs and by playing central roles in biogeochemical cycling of elements. The interactions between phytoplankton and heterotrophic bacteria affect the fitness of both partners. It is becoming increasingly recognized that metabolic exchange determines the nature of such interactions, but the underlying molecular mechanisms remain underexplored. Here, we investigated the molecular and metabolic basis for the bacterial lifestyle switch, from coexistence to pathogenicity, in *Sulfitobacter* D7 during its interaction with *Emiliania huxleyi*, a cosmopolitan bloom-forming phytoplankter. To unravel the bacterial lifestyle switch, we profiled bacterial transcriptomes in response to infochemicals derived from algae in exponential and stationary growth, which induced the *Sulfitobacter* D7 coexistence and pathogenicity lifestyles, respectively. We found that algal dimethylsulfoniopropionate (DMSP) was a pivotal signaling molecule that mediated the transition between the lifestyles. However, the coexisting and pathogenic lifestyles were evident only in the presence of additional algal metabolites. In the pathogenic mode, *Sulfitobacter* D7 upregulated flagellar motility and many transport systems, presumably to maximize assimilation of *E. huxleyi*-derived metabolites released by algal cells upon cell death. Specifically, we discovered that algae-produced benzoate promoted the growth of *Sulfitobacter* D7, and negated the DMSP-inducing lifestyle switch to pathogenicity, demonstrating that benzoate is important for maintaining the coexistence of algae and bacteria. We propose that bacteria can sense the physiological status of the algal host through changes in the metabolic composition, which will determine the bacterial lifestyle during the interactions.

**Significance Statement:** Microorganisms in the marine environment play crucial roles in the regulation of Earth’s climate and elemental cycling. Understanding microbial interactions and the metabolic exchange that drives them is necessary for disentangling the complexity of the marine ecosystem. Here we demonstrate how the opportunistic pathogen *Sulfitobacter* D7 switches its lifestyle from coexistence to pathogenicity in response to metabolites released by *Emiliania huxleyi*, a bloom-forming unicellular alga. By mapping bacterial transcriptional profiles, we show that the algal metabolite dimethylsulfoniopropionate (DMSP), an important signaling molecule in the marine environment, is essential for the bacterial lifestyle switch. However, the activity of DMSP depended on additional algal signals. This work emphasizes how metabolic crosstalk can influence the nature and fate of microbial interactions, which have cascading effects on large-scale oceanic processes.

## Introduction

Half of Earth’s photosynthesis occurs in the marine environment by phytoplankton – photosynthetic single-celled algae (1). Phytoplankton have great ecological importance by forming the basis of marine food webs and influencing biogeochemical cycles. Therefore, the biotic interactions phytoplankton engage in, and the metabolic exchange that govern them, have immense impacts on large-scale biogeochemical processes. Phytoplankton are a main source of organic matter in the marine environment thus fueling the growth and function of heterotrophic bacteria that interact with them through chemical exchange (2, 3). Although the marine environment is characterized by its diluted and turbulent nature, efficient chemical communication takes place in the phycosphere – the diffusive boundary layer that surrounds algal cells, where molecules can accumulate to high concentrations (2, 4). Studies on algae-bacteria interactions revealed that the partners often exchange growth substrates (5, 6), essential vitamins and nutrients (7–9), and infochemicals (molecules that convey information) (10–13). Bacteria have developed mechanisms, such as motility and chemotaxis, for foraging of phytoplankton cells and attachment to their surfaces to maintain close association within the phycosphere (14–21).

Bacteria from the *Rhodobacterales* order, often termed Roseobacters, are found to be associated with phytoplankton (22–29). They are metabolically versatile and specialize on algae-derived substrates that promote interactions with phytoplankton (30). The organosulfur molecule dimethylsulfoniopropionate (DMSP), produced by many phytoplankton species (31), is especially known to mediate Roseobacter-phytoplankton interactions by serving as a carbon and sulfur source, a chemotaxis cue and as an infochemical for the presence of algae (5, 12, 13, 32–36). In mutualistic interactions algae provide organic matter such as sugars, amino acids, sulfonates and polyamines for bacterial growth. In exchange, Roseobacters often produce essential B-vitamins and growth promoting factors such as indole-3-acetic acid (IAA) (5, 12, 37, 38). In recent years cumulating studies that investigated the interactions of phytoplankton and bacteria in co-cultures revealed that some Roseobacters display a lifestyle switch from mutualism to pathogenicity towards the algae. This occurs when the algal host reaches stationary growth and is mediated by infochemicals (6, 7, 13, 39–41). For example, Roseobacters can produce potent algicidal compounds, termed roseobacticides, in response to *p*-coumaric acid, an aromatic lignin breakdown product released by aging algae (11, 34). While this bacterial lifestyle switch, often termed the “Jekyll-and-Hyde” phenotype, seems to be a recurring phenomenon, knowledge about the bacterial behavior in the different modes of interaction and the regulation of such lifestyle switch is still rudimentary.

In the current study we investigated the behavior of the Roseobacter *Sulfitobacter* D7, during interactions with *Emiliania huxleyi*, a cosmopolitan bloom-forming phytoplankter. *E. huxleyi* has a central role in biogeochemical cycling of carbon and sulfur. It produces the climatically active gas dimethyl sulfide and its precursor DMSP, both act as infochemicals during interactions with *E. huxleyi* (13, 42). *Sulfitobacter* sp. are associated with *E. huxleyi* in nature, and *Sulfitobacter* D7 was isolated from a natural *E. huxleyi* bloom (13, 28, 43, 44). Therefore, this ecologically-relevant model provides a tractable system to examine how metabolic exchange regulates the nature of interactions between algae and bacteria. Our previous work revealed that *Sulfitobacter* D7 displays a lifestyle switch, from coexistence to pathogenicity, during its interaction with *E. huxleyi*. We found that algal DMSP, which usually mediates mutualistic interactions, plays a pivotal role by invoking bacterial pathogenicity (13). Many studies investigated the genes related to DMSP uptake and catabolism (45–49), but the regulation and molecular basis of DMSP-responsive genes and how these affect bacterial lifestyle and behavior during interactions with algae are yet to be explored. We performed a transcriptome experiment that allowed us to elucidate the bacterial response to algal infochemicals, and to characterize DMSP-responsive and pathogenicity-related genes. We revealed the signaling role of DMSP that leads to systemic remodeling of *Sulfitobacter* D7 gene expression, but only in the presence of additional algal metabolites. Overall, we unravel the transcriptional signature of the shift from coexistence to pathogenic bacterial lifestyles during interactions with their algal host and provide insights into the ecological context of these modes of interactions.

## Results

### *E. huxleyi*-derived exudates induce remodeling of *Sulfitobacter* D7 transcriptome

The interaction between *Sulfitobacter* D7 and *E. huxleyi* displays two distinct phases (Fig. 1a). Initially, there is a coexisting phase in which the alga grows exponentially and the bacterium grows as well. The interaction shifts to pathogenic when the virulence of *Sulfitobacter* D7 towards *E. huxleyi* is invoked upon exposure to high concentrations of algal DMSP, which occurs when the algae reach stationary growth or when DMSP is applied exogenously to algae in exponential growth (Fig. 1a) (13). We aimed to unravel the response of *Sulfitobacter* D7 to the pathogenicity-inducing compound, DMSP, and to different algae-derived infochemicals that affect the lifestyle of the bacterium. We grew *Sulfitobacter* D7 in conditioned media (CM) derived from algal cultures at exponential and stationary phase (Exp-CM and Stat-CM, respectively), in which DMSP concentration is low and high, respectively (13) (Fig. 1b, Table S1). This enabled us to dissect the interaction with *E. huxleyi* into its different phases, i.e., Exp-CM represents the coexisting phase, and Stat-CM represents the pathogenic phase. An additional pathogenicity-inducing treatment was Exp-CM supplemented with 100 μM DMSP (herein Exp-CM+DMSP). This condition mimicked co-cultures to which we added DMSP exogenously and thus induced *Sulfitobacter* D7 pathogenicity, which lead to death of exponentially growing *E. huxleyi*. In order to reveal the bacterial transcriptional response to algal exudates, we harvested bacterial cells after 24 h of growth in the different CM treatments and performed RNAseq analysis, using a modified protocol based on Avraham *et al*., 2016 (50) (Fig. 1b, Table S1). We aimed to identify pathogenicity-related genes by comparing *Sulfitobacter* D7 gene expression profiles in the pathogenicity-inducing media to the coexistence medium. We also aimed to identify bacterial genes that are specifically responsive to DMSP, and are not affected by other algae-derived factors. Therefore, we grew *Sulfitobacter* D7 in defined minimal medium (MM), which lacks algal metabolites, supplemented it with 100 μM DMSP (herein MM+DMSP), and examined the transcriptional response. This experimental design allowed us to expand our understanding on the bacterial response to DMSP, algal infochemicals and which of these are essential for coexistence and pathogenicity.

**Fig. 1.**
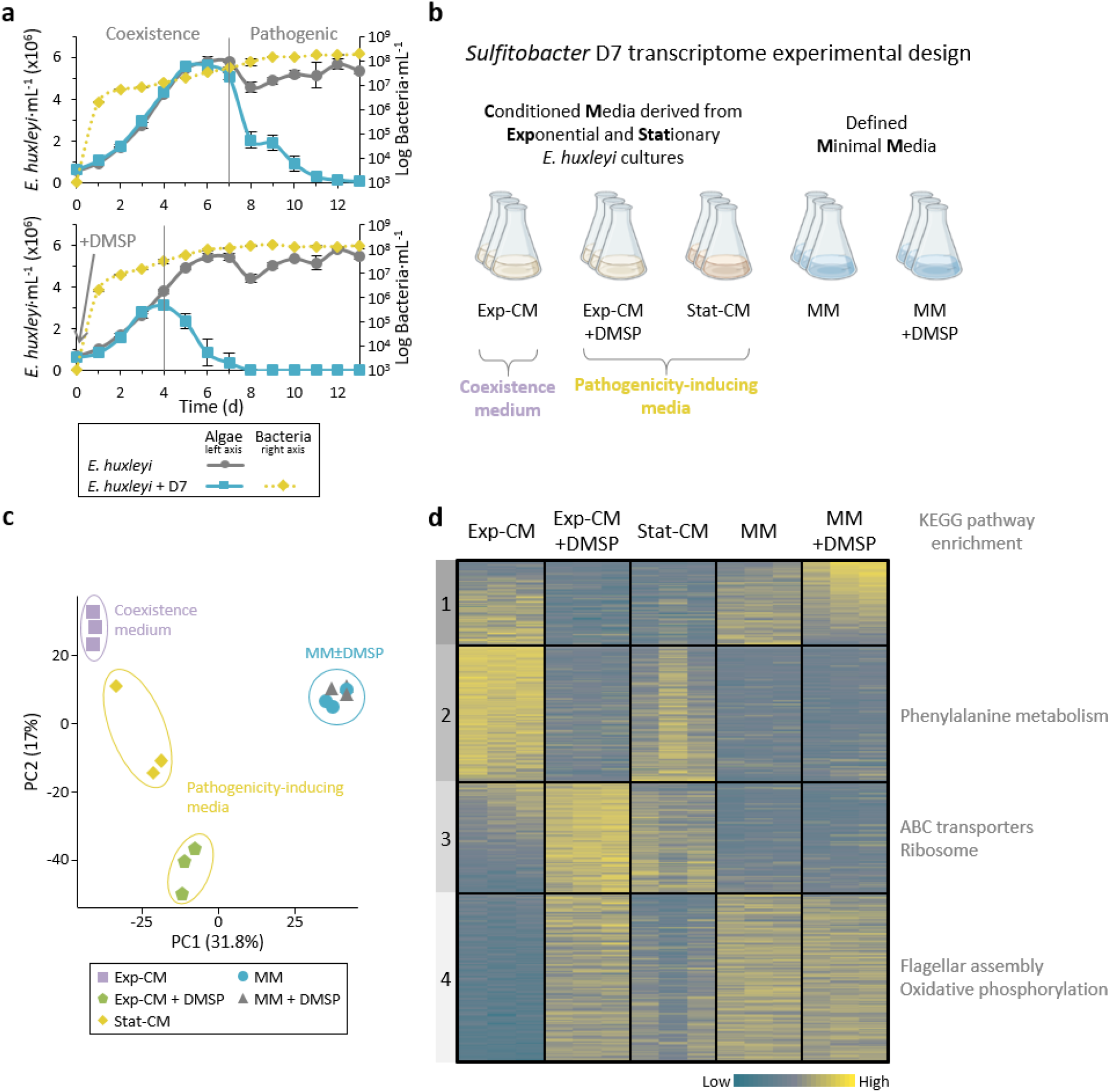
Transcriptional profiling of *Sulfitobacter* D7 in response to DMSP and additional *E. huxleyi* infochemicals reveals the signaling role of DMSP. (a) Time course of *E. huxleyi* (strain CCMP379) and bacterial abundance (full and dashed lines, left and right axes, respectively) in algal mono-cultures or during co-culturing with *Sulfitobacter* D7. Top panel: Co-cultures display two phases with distinct bacterial lifestyles: coexistence and pathogenicity. Bottom panel: DMSP was added at day 0 to a final concentration of 100 μM. Results represent average ± SD (n = 3). Adapted from Barak-Gavish *et al*., 2018 (13) (b) Design of *Sulfitobacter* D7 transcriptome experiment aiming to explore gene expression profiles in response to *E. huxleyi*-derived media and in response to DMSP. Growth media consisted of conditioned media (CM) derived from *E. huxleyi* at exponential or stationary growth phase (Exp-CM and Stat-CM, respectively), and an additional treatment in which 100 μM of DMSP was added (Exp-CM+DMSP). These media differentially induce the coexistence and pathogenicity lifestyles of *Sulfitobacter* D7. In order to identify DMSP-responsive genes we inoculated *Sulfitobacter* D7 in defined minimal media (MM), lacking *E. huxleyi*-derived exudates, without and with 100 μM DMSP (MM and MM+DMSP, respectively). *Sulfitobacter* D7 was inoculated into each media and harvested for RNA profiling after 24 h of growth. Initial conditions of the media and bacterial growth are elaborated in Table S1. (c) Principle component analysis of *Sulfitobacter* D7 detected genes in all treatments (2588 genes). Triplicates of each treatment are shown. We observed a clear separation between CM treatments and the MM treatments. (d) Heatmap of gene expression of all differentially expressed (DE) genes in the comparisons Exp-CM+DMSP vs. Exp-CM, Stat-CM vs. Exp-CM and MM+DMSP vs. MM (1179 genes). DE genes were defined as genes with |fold change|>2 and adjusted *P*-value ≤ 0.05. Clusters were determined based on k-means analysis. Significant functional enrichment in each cluster, based on KEGG Pathways, are denoted. Each row represents one gene and the color intensity corresponds to the standardized expression across all samples (triplicates of each treatment are shown). Expression values are scaled by row. Genes in cluster 1 are ordered based on the mean expression values in the MM+DMSP treatment. Genes in cluster 2-4 are ordered based on the mean expression values in the Exp-CM treatment.

Among the 3803 genes in *Sulfitobacter* D7 genome, we detected 2588 genes (Table S2). Principle component analysis (PCA), based on the expression profile of all the detected genes, showed a clear separation between the different CM treatments, while the MM ± DMSP samples clustered together (Fig. 1c). Pearson correlation between all samples indicated high correlation between the triplicates of each treatment and hierarchical clustering showed that MM samples cluster separately from the CM samples (Fig. S1). Among the CM samples, Stat-CM and Exp-CM+DMSP were closer and had high correlation to each other compared to Exp-CM (Fig. S1). This suggests that the pathogenicity-inducing impact of the former media is mediated by a unique set of expressed genes, distinct from that induced by the coexistence medium.

In order to identify the pathogenicity- and DMSP-related genes we examined the genes that were differentially expressed (DE) in the following comparisons: Exp-CM+DMSP vs. Exp-CM, Stat-CM vs. Exp-CM and MM+DMSP vs. MM. We defined significantly DE genes as |fold change|>2 and adjusted *P*-value ≤ 0.05. The DE genes were separated into 4 clusters, based on k-means clustering, and we assessed the enrichment in KEGG pathways in each cluster (Fig. 1d). Cluster 1 contained genes that were responsive to DMSP, namely DE in Exp-CM±DMSP and in MM±DMSP comparisons. Interestingly, the expression pattern of cluster 1 genes in response to DMSP was largely different: in Exp-CM, DMSP lead to downregulation, while in MM it lead to upregulation. The differential effect of DMSP in Exp-CM and MM is also visualized in the PCA and is evident by the number of DE genes in the comparisons: 968 genes were DE between Exp-CM vs. Exp-CM+DMSP, while only 170 genes were DE between MM vs. MM+DMSP (Table S3). Since the metabolic composition of these two media was profoundly different, it suggests that the effect of DMSP signaling depends also on the chemical environment. Namely, in different chemical contexts DMSP will affect *Sulfitobacter* D7 gene expression in a different way.

Cluster 2 contained genes that were highly expressed in the coexistence medium, Exp-CM, compared to the pathogenicity-inducing media, therefore, we consider it as coexistence-related. This cluster was enriched with genes related to the phenylalanine metabolism pathway, and specifically to phenylacetic acid (PAA) degradation (Table S4). PAA is a phytohormone that can potentially promote algal growth (51), and PAA metabolism by bacteria is related to production of secondary metabolites that can affect algae (52, 53). Clusters 3 and 4 contained genes that were mainly upregulated in Exp-CM+DMSP and Stat-CM compared to Exp-CM, and we consider these as pathogenicity-related clusters. Cluster 4 also contained genes that were expressed in MM±DMSP. Cluster 3 was enriched with genes encoding for ABC transporters and ribosomal proteins, and cluster 4 was enriched with flagellar genes and genes related to oxidative phosphorylation (Fig. 1d, Table S4). The enrichment in genes encoding for ribosomal proteins and an F-type ATPase, related to oxidative phosphorylation, in the pathogenicity-related clusters suggests that during the pathogenic lifestyle *Sulfitobacter* D7 is more metabolically active than in the coexistence lifestyle (Table S4). Overall, this transcriptome experimental setup enabled us to capture the gene expression of *Sulfitobacter* D7 in coexistence and pathogenicity modes. Moreover, it demonstrated the pivotal role of DMSP in regulating gene expression in a context-dependent manner. In *E. huxleyi*-derived CM, DMSP led to major changes in *Sulfitobacter* D7 transcriptome, while in MM, which lacked additional algal metabolites, DMSP had a minor effect on gene expression. We therefore suggest that the additional algal factors act in concert with DMSP and are required for the expression of coexistence-and pathogenicity-related genes.

### The pathogenic lifestyle of *Sulfitobacter* D7 includes upregulation of flagellar genes and increased motility

The enrichment in flagellar genes in cluster 4 suggests that flagellar motility may be involved in the pathogenic lifestyle of *Sulfitobacter* D7. We examined the expression of all the genes necessary for flagellar assembly, which are organized in an operon-like structure in *Sulfitobacter* D7 genome (Fig. 2a). Most of the flagellar genes were significantly upregulated in pathogenicity-inducing media compared to the coexistence medium (Fig. 2a). This includes the majority of the genes encoding for the flagellar hook and basal body (Fig. 2a and Table S5). The genes FliC, FliM, FlgC, FlgB and FliI were not DE but were highly expressed in all treatments (Table S5). The genes FlhB, FliR, FlhA and FliQ were not sufficiently detected in our analysis. In MM, however, there were no significant changes in expression of flagellar genes in response to DMSP, although the overall expression was higher than in the CM samples (Fig. 2a, Table S5).

**Fig. 2.**
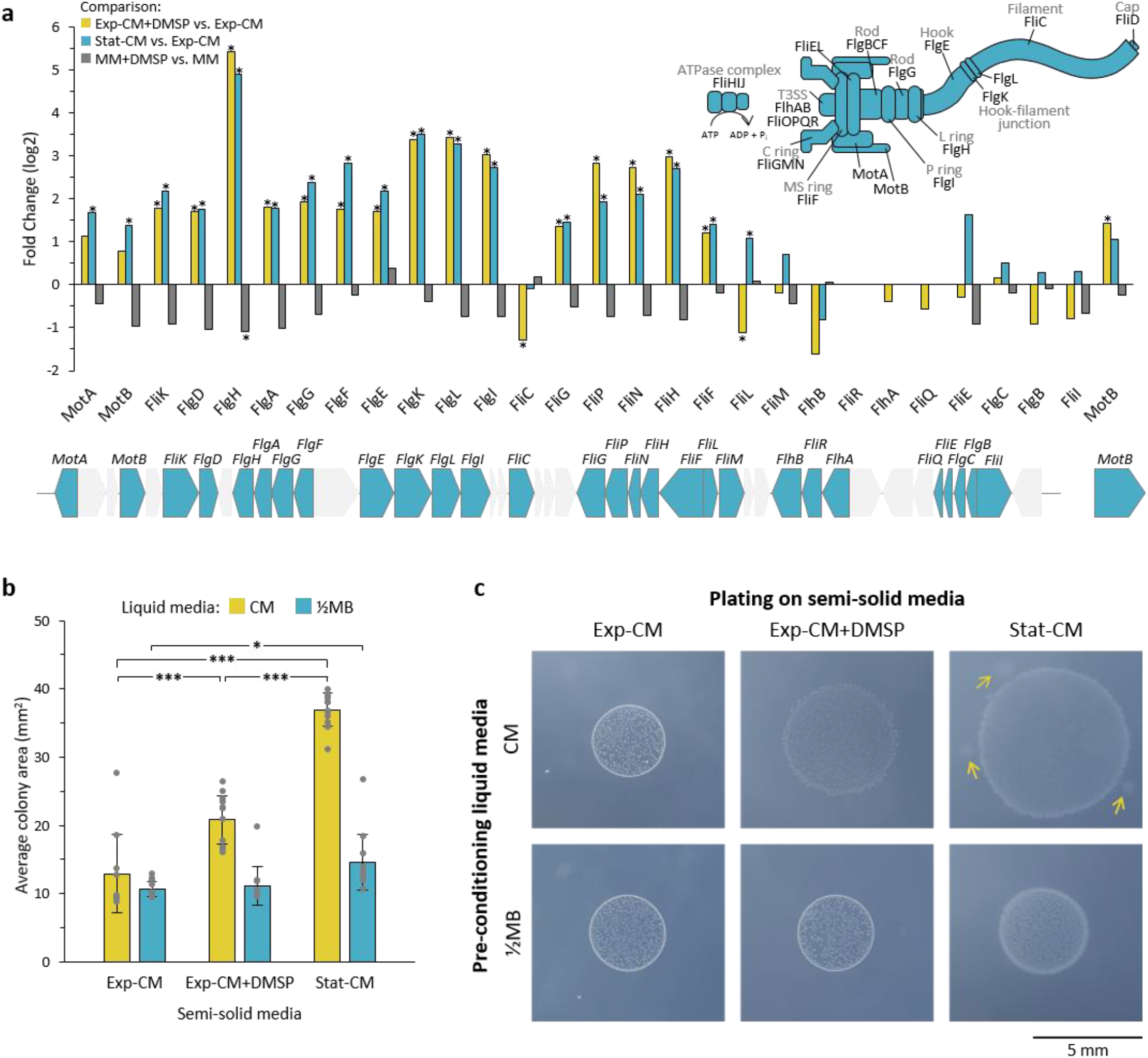
Induction of flagellar genes and increased motility in pathogenicity-inducing media. (a) Fold change of flagellar gene expression in the comparisons: Exp-CM+DMSP vs. Exp-CM (yellow), Stat-CM vs. Exp-CM (blue) and MM+DMSP vs. MM (grey). Genes marked with * are significantly DE. The flagellar genes are encoded on the *Sulfitobacter* D7 chromosome in an operon-like structure, as shown below the graph. Genes in grey are not related to the flagellum. The function of each gene is indicated in the flagellum assembly scheme on the top-right. Expression values are presented in Table S5. (b) Bacterial motility inferred by the colony area of *Sulfitobacter* D7 plated on semi-solid agar media (0.3% agarose). Bacteria were pre-conditioned in liquid CM (Exp-CM, Exp-CM+DMSP and Stat-CM) for 24 h and plated on the corresponding semi-solid CM plates (yellow bars). For control, bacteria were pre-conditioned on ½MB and plated on semi-solid CM plates (blue bars). Colony area was determined after 6 days of growth. Results represent average ± SD (n=10-12 colonies per treatment). * *P*-value<0.05, *** *P*-value<0.0001. (c) Representative bacterial colonies from each treatment showing the difference in colony area and morphology. The arrows depict bacterial motility extensions from the core colony. The extensions were not included in the colony area measurements.

To assess the involvement of motility in the behavioral switch of *Sulfitobacter* D7 and to validate the expression patterns of flagellar genes, we performed a functional bacterial motility assay in response to *E. huxleyi*-derived metabolites. We examined the colony expansion of *Sulfitobacter* D7 in semi-solid agar plates composed of Exp-CM, Exp-CM+DMSP or Stat-CM. We first pre-conditioned *Sulfitobacter* D7 in the respective liquid CM for 24 h, in order to induce the appropriate expression of flagellar genes. *Sulfitobacter* D7 plated on pathogenicity-inducing semi-solid media showed higher colony expansion area than in the coexistence medium, indicating increased motility in these conditions (Fig. 2 b-c). The average colony area in semi-solid Stat-CM and Exp-CM+DMSP was 37 and 20.8 mm^2^, respectively, while in Exp-CM it was only 13 mm^2^ (Fig. 2b). Moreover, the morphology of the colonies was different; the colony edges in Stat-CM were smeared and there were bacterial motility extensions from the core colony, indicating bacterial migration in the semi-solid agar (Fig. 2c). The smeared edges were also evident in Exp-CM+DMSP, but to a lesser extent. These results validated the expression patterns of flagellar genes in each CM. Interestingly, *Sulfitobacter* D7 that was pre-grown in liquid marine broth (½MB), and was therefore not pre-exposed to *E. huxleyi* infochemicals, and subsequently plated on the three semi-solid CM did not show major differences in the average colony area. This strongly indicates that *Sulfitobacter* D7 grown in liquid pathogenicity-inducing media were pre-conditioned for motility by upregulating the expression of flagellar genes compared to the coexistence medium. Nevertheless, colonies plated on Stat-CM were significantly larger and showed the smeared edges morphology, implying that in semi-solid Stat-CM there was also induction of motility. Taken together, high expression of flagellar genes in pathogenicity-inducing media, along with the observation that bacteria are indeed more motile in these conditions, indicate that flagella-driven motility may be involved in the pathogenic lifestyle of *Sulfitobacter* D7 during interaction with *E. huxleyi*.

### DMSP and *E. huxleyi*-derived metabolites modulate the expression of *Sulfitobacter* D7 transport genes

The enrichment in ABC transporters in cluster 3 suggests that nutrient uptake by *Sulfitobacter* D7 is prominent during the interaction with *E. huxleyi*. We examined the expression of all 493 transport genes in *Sulfitobacter* D7 genome. We found that transporters for energy-rich organic compounds were expressed in CM treatments (Fig. 3, Table S6). This includes transporters for amino acids and peptides, carbohydrates and sugars, organic sulfur and nitrogen compounds, as well as for inorganic nutrients and metals. Examination of bacterial transport genes was shown to serve as a sensitive readout for estimating which metabolites reside in the media and are taken up by bacteria (54). Therefore, the expression of transporters implies that the CM contained *E. huxleyi*-derived metabolites that *Sulfitobacter* D7 can benefit from during growth in CM and during the interaction with the alga. Such metabolites include branched-chain amino acids, sugars, C4 carbohydrates and DMSP, which are known to be produced by *E. huxleyi* (55, 56) (Fig. 3).

**Fig. 3.**
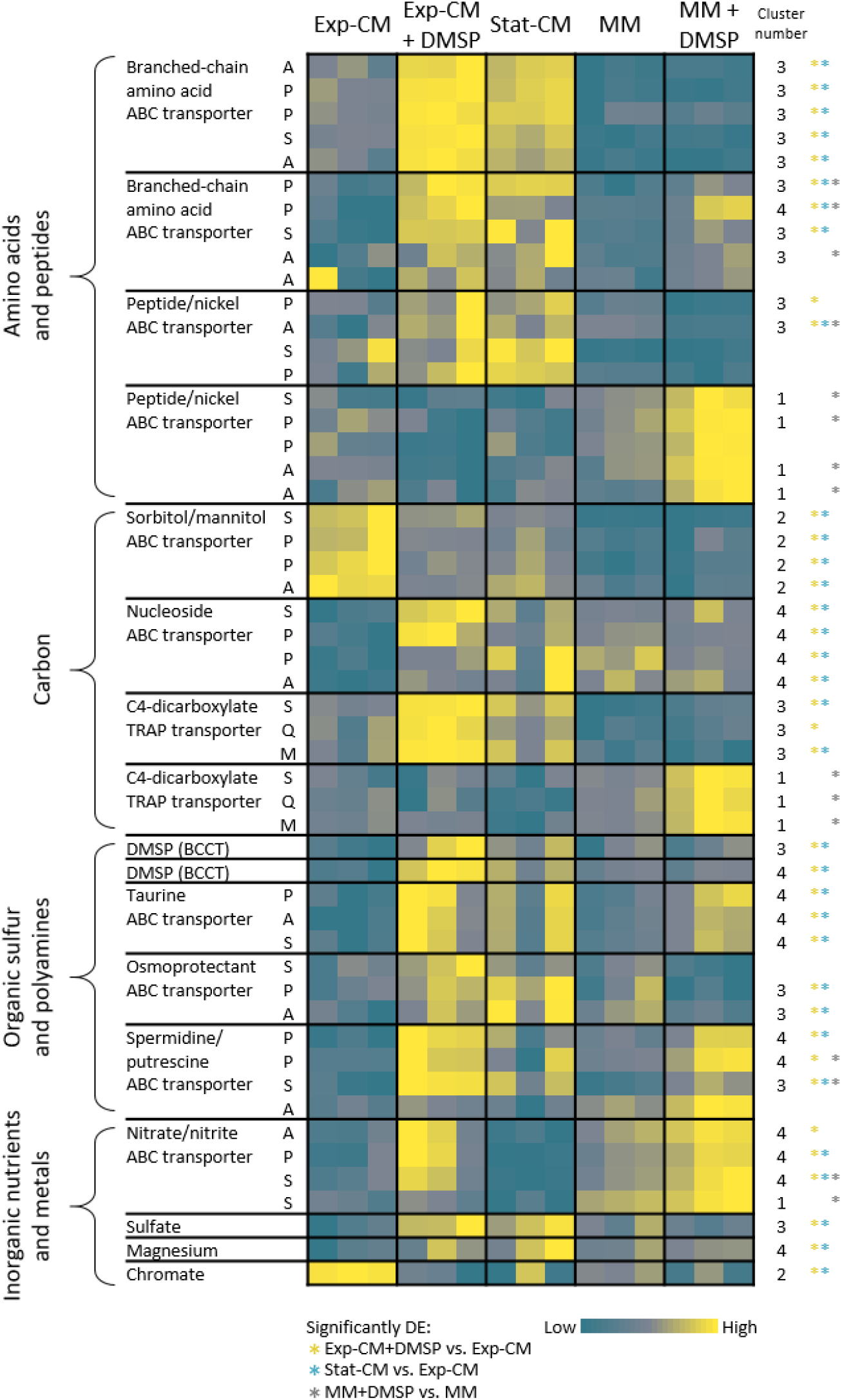
Remodeling of *Sulfitobacter* D7 transport systems in response to DMSP and *E. huxleyi*-derived metabolites. Heatmap of gene expression of representative transport genes for various metabolite classes. Each row represents one gene and the blocks represent a complete transport system in which at least two genes were DE in the comparisons indicated on the bottom. The column “cluster number” corresponds to the heatmap-cluster in Fig. 1d in which the gene is found. Colored * denote in which comparison the gene was significantly DE. Color intensity correspond to the standardized expression across all samples (triplicates of each treatment are shown). Expression values are scaled by row. Expression and fold-change values are presented in Table S6. ABC, ATP-binding cassette; TRAP, tripartite ATP-independent periplasmic; BCCT, betaine/Carnitine/Choline Transporter. The letters corresponds to the transport system components: A, ATP-binding; S, substrate-binding; P, permease; Q, dctQ subunit; M, dctM subunit.

Numerous transport genes were DE in the pairwise comparisons of the different treatments (Fig. 3, Fig. S2 and Table S6). *Sulfitobacter* D7 grown in Stat-CM had a similar expression profile of transport genes to that of Exp-CM+DMSP (Fig. 3, Fig. S2 and Table S6). Therefore, *Sulfitobacter* D7 grown in the pathogenicity-inducing media was indeed in a unique transcriptional and metabolic state compared to the coexistence medium. Many transport genes were mostly upregulated in Exp-CM+DMSP compared to Exp-CM: 99 genes, which constitute ∼20% of *Sulfitobacter* D7 transport genes (Fig. 3, Fig. S2). An additional 39 genes were downregulated in response to DMSP addition to Exp-CM. Interestingly, also in MM+DMSP 42 transport genes were upregulated compared to MM, and 14 were downregulated. Namely, in both Exp-CM and MM, DMSP induced remodeling of the transporter repertoire. The fact that the addition of a single metabolite, i.e. DMSP, lead to DE of a multitude of transporters for various metabolite classes demonstrates the signaling role of DMSP. When we examined the amount of DE transport genes that were shared between the comparisons of the DMSP-added samples, we found only 8 genes were differentially expressed in a similar manner (Fig. S2). Namely, DMSP lead to a shift in transporter gene expression in both media but the identity of the DE transporters was unique for each media. This strengthens our hypothesis that the DMSP signal affects bacterial gene expression but the activation of coexistence and pathogenic transcriptional profile depends also on additional algal factors.

The differential effect of DMSP in Exp-CM and MM was especially notable in the expression of the DMSP transporters (BCCT, betaine-carnitine-choline transporter), which were significantly upregulated in Stat-CM and Exp-CM+DMSP, where DMSP was present at high concentrations, compared to Exp-CM. However, the expression of these transporters was not affected by the addition of DMSP in the MM±DMSP treatments. This was the same for the *DmdA* gene that encodes for the enzyme responsible for the first step of DMSP breakdown (49) (Table S7). *DmdA* was barely expressed in MM+DMSP, albeit DMSP was present in high concentrations. Therefore, DMSP uptake and metabolism were prominent in *E. huxleyi*-derived CM, which contain additional algal factors that are not present in MM. Taken together, it seems that DMSP has a strong signaling role, however, additional algal components are required to induce the coexistence-to pathogenicity-related gene expression in *Sulfitobacter* D7.

### Algal benzoate is a key metabolite for *E. huxleyi*-*Sulfitobacter* D7 coexistence

We searched in the *Sulfitobacter* D7 transcriptomic response to *E. huxleyi*-derived metabolites for evidence of involvement of additional algal factors, other than DMSP, in the regulation of the lifestyle switch from coexistence to pathogenicity. We revealed a plasmid-encoded degradation pathway of the aromatic compound benzoate that was highly expressed in CM samples (Fig. 4a-b, Table S8). Aromatic compound degradation is a common metabolic feature in Roseobacters (57, 58). The metabolic intermediates of benzoate catabolism can be directed to β-ketoadipate, which is subsequently metabolized to form the TCA precursors acetyl-CoA and succinate (59). There are two pathways to metabolize benzoate to β-ketoadipate, through catheol (*Ben* genes) and through 4-hydroxybenzoate and protocatecuate (*BphA* and *PobA* genes) (Fig. S3). An additional benzoate degradation pathway is through benzoyl-CoA (*Box* genes) (Fig. S3) (60). *Sulfitobacter* D7 harbors the pathway through catechol (Fig. 4a). The *Ben* genes are organized in an operon-like structure adjacent to a transcription factor *BenM*, which is known to regulate the expression of *BenABCD* and *CatAB* (Fig 4b) (61, 62). All the genes in this benzoate-degradation operon were expressed in CM treatments. Interestingly, the transporter of benzoate, which is encoded by a chromosomal gene, was also expressed in all CM, therefore it is not affected by the DMSP signal, as was observed for other transport systems (Fig. 3b). This suggest that *Sulfitobacter* D7 can assimilate and perceive algae-derived benzoate as a growth factor or signal regardless of the concentration of DMSP and may therefore be important in the initial coexistence phase.

**Fig. 4.**
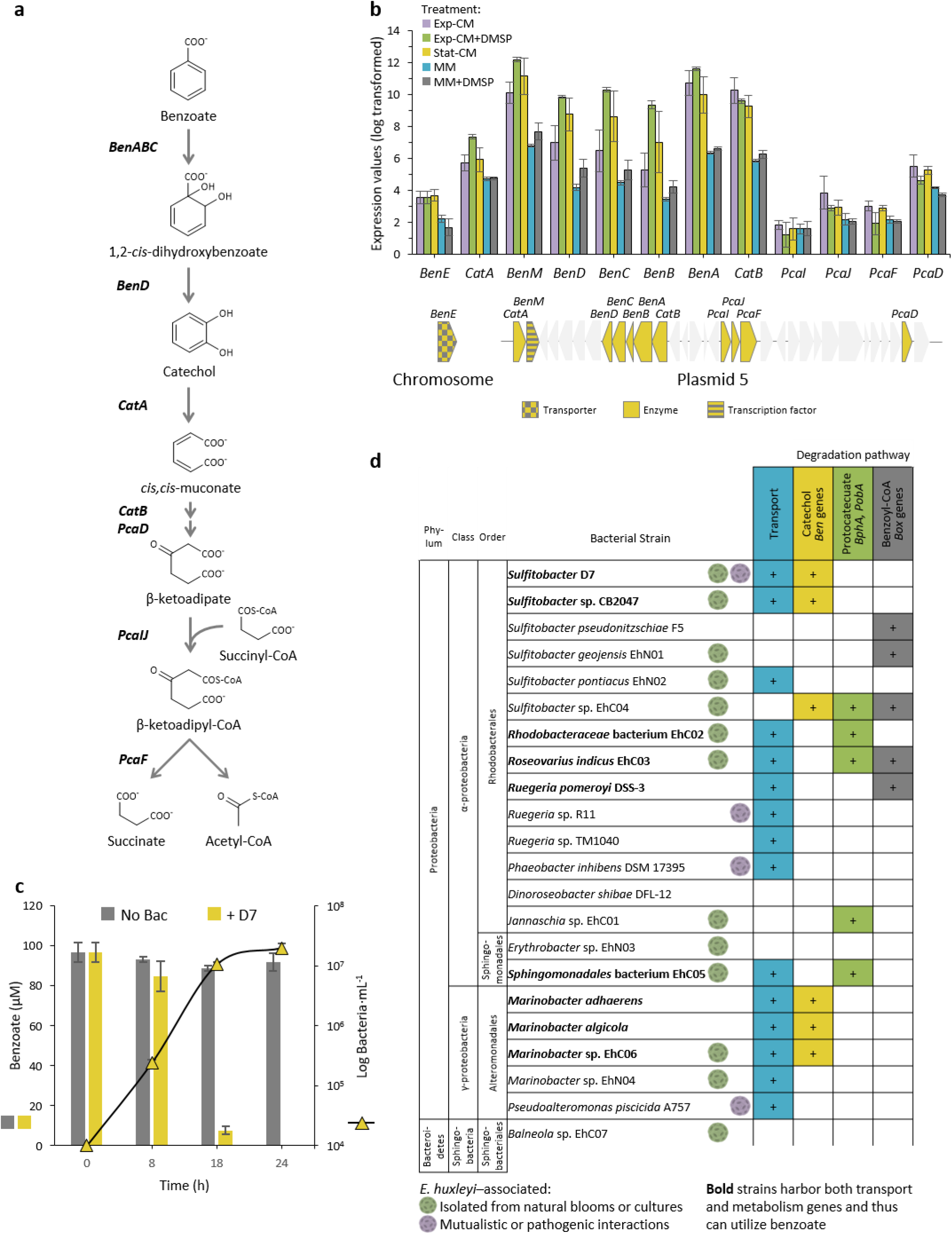
*Sulfitobacter* D7 encodes for a benzoate degradation pathway required for the metabolic exchange with *E. huxleyi*. (a) The benzoate degradation pathway in *Sulfitobacter* D7. The genes that encode for the enzymes mediating the subsequent transformations of benzoate to succinate and acetyl-CoA are denoted in bold. (b) Expression values of benzoate-related genes, which are encoded on *Sulfitobacter* D7 plasmid 5 in an operon-like structure, as indicated below the graph. The benzoate transporter, *BenE*, is encoded on the chromosome. Genes in grey are not related to benzoate. Results represent average ± SD (n = 3). (c) Benzoate concentration (bars, left axis) and bacterial growth (triangles, right axis) in MM supplemented with 100 μM benzoate, as a sole carbon source, without inoculation (grey) and upon inoculation of *Sulfitobacter* D7 (yellow). Results represent average ± SD (n = 3). *P*-value<0.0001 for the difference in benzoate concentration between the “No bac” and “+D7” treatments. No bacterial growth was observed in un-inoculated MM. (d) Presence of benzoate transport and degradation genes in genomes of phytoplankton-associated bacteria. *E. huxleyi*-associated bacteria are denoted. Colored tiles depict the presence of the genes. Bacterial strains highlighted in bold possess genes for both transport and degradation of benzoate. Bacterial benzoate degradation pathways are elaborated in Fig. S3. The full data of the presence of all query genes in the genomes of the bacteria is presented in Table S9. The query genes are listed in Table S10. *BenABC*, benzoate 1,2-dioxygenase subunit alpha, beta and reductase component, respectively; *BenD, cis*-1,2-dihydroxybenzoate dehydrogenase; *CatA*, catechol 1,2-dioxygenase; *CatB*, muconate cycloisomerase; *PcaD*, 3-oxoadipate enol-lactonase; *PcaIJ*, 3-oxoadipate CoA-transferase, alpha and beta subunits, respectively; *PcaF*, 3-oxoadipyl-CoA thiolase.

To test if *Sulfitobacter* D7 can grow and metabolize benzoate we inoculated the bacterium in MM supplemented with 100 μM benzoate as a sole carbon source. *Sulfitobacter* D7 grew by 3 orders of magnitude within 24h and consumed benzoate to an undetectable level, while the concentration of benzoate in the non-inoculated medium did not change significantly (Fig. 4c). We also detected benzoate in the media of exponentially growing *E. huxleyi* cultures (Fig. S4). This suggests that *Sulfitobacter* D7 can grow on benzoate and benefit from this metabolite during interactions with *E. huxleyi*.

Bacterial degradation of various aromatic compounds is mostly directed to the β-ketoadipate pathway and eventually to the TCA cycle (59). While this pathway seems to exist in many Roseobacters, the direct degradation of benzoate is limited to only few species (30, 57). We examined the prevalence of benzoate degradation and transport genes among phytoplankton-associated bacteria. Specifically, we searched for benzoate transporters and genes that directly metabolize benzoate through one of the three possible pathways (Fig. S3, Table S9 and S10). We found that in addition to *Sulfitobacter* D7 another *Sulfitobacter* sp., CB2047, which was also isolated from an *E. huxleyi* bloom (44), was able to utilize benzoate, namely, its genome encodes for both degradation and transport genes (Fig. 3d). We found indications for benzoate utilization in the genomes of two additional *Roseobacter*s, *Rhodobacteraceae* bacterium EhC02 and *Roseovarius indicus* EhC03, as well as *Sphingomonadales* bacterium EhC05, all isolated from *E. huxleyi* cultures (63) (Fig. 3d). This was also evident in the genomes of several *Marinobacter*s, a genus known to be associated with *E. huxleyi* cultures (63–65). *Ruegeria pomeroyi* DSS-3 was the only bacterial strain included in our analysis, which is not directly associated to *E. huxleyi* but showed benzoate utilization, consistent with previous observations (30). This suggests that benzoate produced by *E. huxleyi* can mediate interactions of the alga with other bacteria that consume and benefit from this metabolite.

We next examined the impact of benzoate on the lifestyle of *Sulfitobacter* D7 during interactions with *E. huxleyi*. We followed the dynamics of a co-culture of *Sulfitobacter* D7 and *E. huxleyi* (strain CCMP2090) in which the bacterial lifestyle switch, from coexistence to pathogenic, does not naturally occur (Fig. 5a-b). This is attributed to the low concentrations of DMSP in the media of this specific *E. huxleyi* strain (13). Indeed, bacterial pathogenicity was induced only when we added external DMSP, leading to the decline of *E. huxleyi* abundance (Fig. 5b). These results are consistent with our previous findings (13). When we added benzoate we did not observe any change in the dynamics of algal growth, but bacterial growth was enhanced by 10-fold at day 3, compared to the non-supplemented and DMSP-supplemented co-cultures (Fig. 5c). This goes along with the observation of benzoate being a remarkable metabolite for *Sulfitobacter* D7 growth (Fig. 4C). When we applied both benzoate and DMSP, bacterial growth was enhanced and algal growth was not compromised although *Sulfitobacter* D7 and DMSP were present in high concentrations. Intriguingly, the presence of benzoate negated the pathogenicity-inducing effect of DMSP on *Sulfitobacter* D7. Moreover, bacterial abundance in both benzoate-added treatments was higher than in the pathogenicity-inducing DMSP treatment, which demonstrates that the onset of pathogenicity is decoupled from bacterial density. This suggests that benzoate is important to maintain *E. huxleyi*-*Sulfitobacter* D7 coexistence and prevents the onset of bacterial pathogenicity. This observation strengthens our previous conclusion that DMSP signaling in *Sulfitobacter* D7 depends on additional algae-derived metabolites which affect the bacterial lifestyle switch during interactions with *E. huxleyi*.

**Fig. 5.**
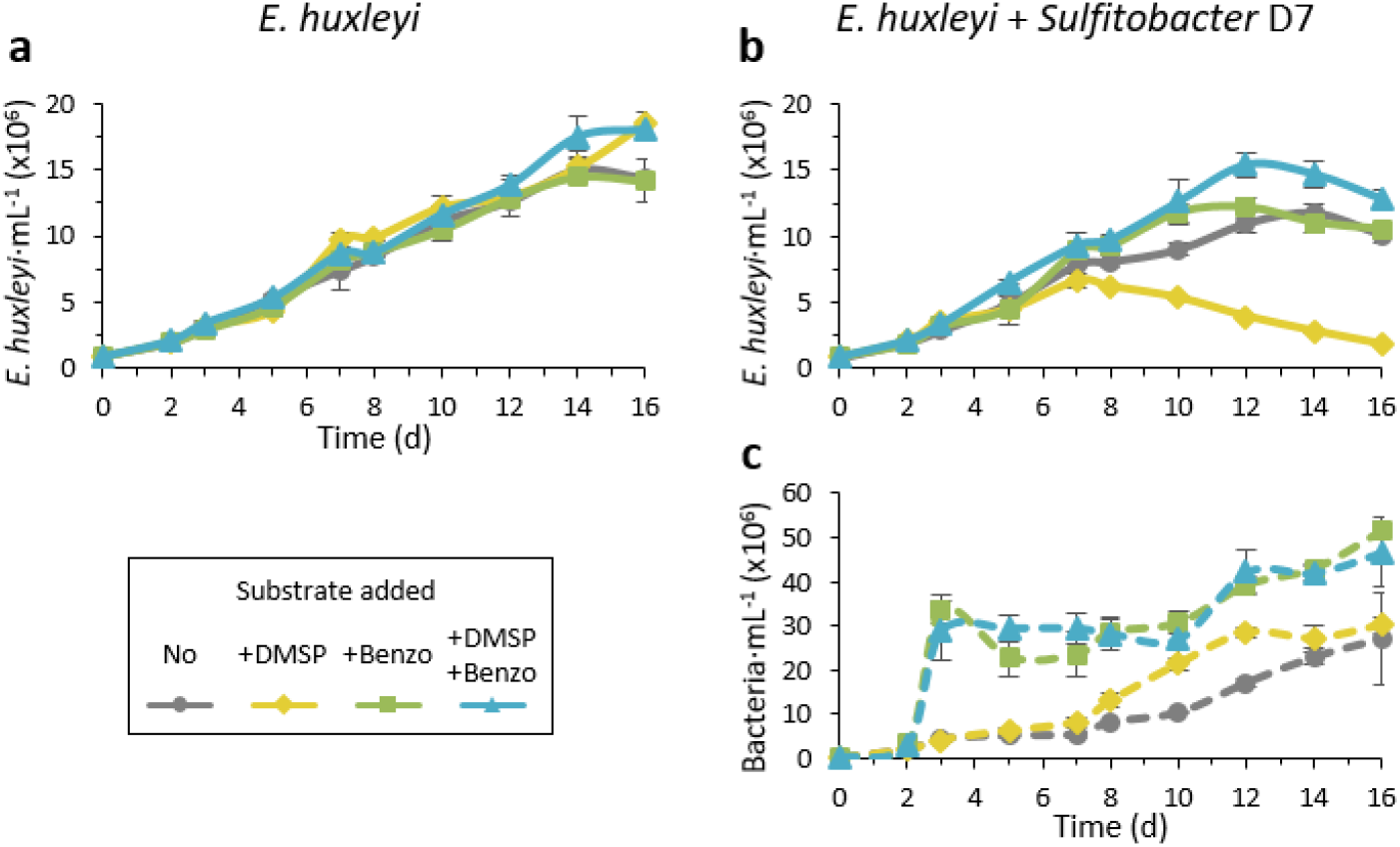
Benzoate is a key metabolite for maintaining *E. huxleyi*-*Sulfitobacter* D7 coexistence. Time course of *E. huxleyi* (strain CCMP2090) and bacterial abundance (smooth and dashed lines, respectively) in algal mono-cultures (a) or during co-culturing with *Sulfitobacter* D7 (b-c). No bacterial growth was observed in algal mono-cultures. Cultures were supplemented at day 0 with 100 μM of DMSP (yellow), benzoate (benzo, green), DMSP and benzoate (blue) or none (grey). The presence of benzoate negated the pathogenicity-inducing effect of DMSP. Results represent average ± SD (n = 3). *P*-value<0.0001 for the difference in *E. huxleyi* growth in the treatment of “+D7+DMSP” compared to all other treatments. *P*-value<0.05 for the differences in bacterial growth in the treatments “+D7+benzoate” and “+D7+Benzoate+DMSP” compared to only “+D7”. *P*-values for all comparisons are listed in Table S11.

## Discussion

### Signaling role of DMSP and other algal infochemicals in the lifestyle switch of *Sulfitobacter* D7

In this study, we aimed to unravel the molecular basis for the lifestyle switch from coexistence to pathogenicity in *Sulfitobacter* D7 during interactions with the bloom-forming algae *E. huxleyi*. We substantiated the signaling role of algal DMSP that mediates the shift towards pathogenicity by mapping the transcriptional profiles of *Sulfitobacter* D7 in response to DMSP and other algal infochemicals. However, DMSP signaling in media that lacked *E. huxleyi*-derived metabolites (i.e., MM±DMSP) had a different effect on *Sulfitobacter* D7 transcriptome. We propose that the signaling role of DMSP that mediates the coexistence to pathogenicity lifestyle switch in *Sulfitobacter* D7 depends on other infochemicals produced by *E. huxleyi*. DMSP is a ubiquitous infochemical produced by many phytoplankton species as well as some bacteria (66), making it a prevalent signaling molecule that mediates microbial interactions in the marine environment. Therefore it is likely that other algal metabolites are involved in the recognition of the specific phytoplankter host by bacteria, thus ensuring specificity in DMSP signaling during interactions. In complex environments, where many microbial species are present simultaneously, such a mechanism can ensure that bacteria will invest in altering gene expression and metabolic remodeling only when the right algal partners are present.

We revealed that the alga-derived aromatic compound benzoate plays a pivotal role in *Sulfitobacter* D7-*E. huxleyi* interaction by maintaining the coexistence, even when DMSP was present at high concentrations (Fig. 5). Benzoate also acts as an efficient bacterial growth factor serving as a carbon source. These observations provide a possible explanation for the switch in bacterial behavior from coexistence to pathogenicity. During the interaction, *E. huxleyi* provides benzoate and other growth substrates to *Sulfitobacter* D7, which uptakes and consumes them (Fig. 6). We propose that as long as *Sulfitobacter* D7 benefits from the interaction with *E. huxleyi* by receiving beneficial growth substrates it will maintain in a coexisting lifestyle. When the growth substrates provided by the alga are less beneficial, the opportunistic pathogen will switch to killing the algal host, which will in turn lead to a surge of intracellular *E. huxleyi*-derived metabolites that *Sulfitobacter* D7 can benefit from (Fig. 6). Studies on phytoplankton exudation of organic matter demonstrated that algae release more organic matter in stationary growth, but the chemical composition is different than that of exponential growth (67, 68). In nutrient limiting conditions, which often occurs in stationary phase, the organic matter exuded by phytoplankton is less favorable for bacterial uptake and consumption for growth (69). In such a chemical context, high concentrations of algae-derived infochemicals, e.g. DMSP, can be perceived by bacteria and signal that the physiological state of the algal host is deteriorating. Namely, by sensing the change in the metabolic composition *Sulfitobacter* D7 executes its pathogenicity against a compromised *E. huxleyi* population. Therefore, the initial coexistence phase is a prerequisite for the onset of bacterial pathogenicity.

**Fig. 6.**
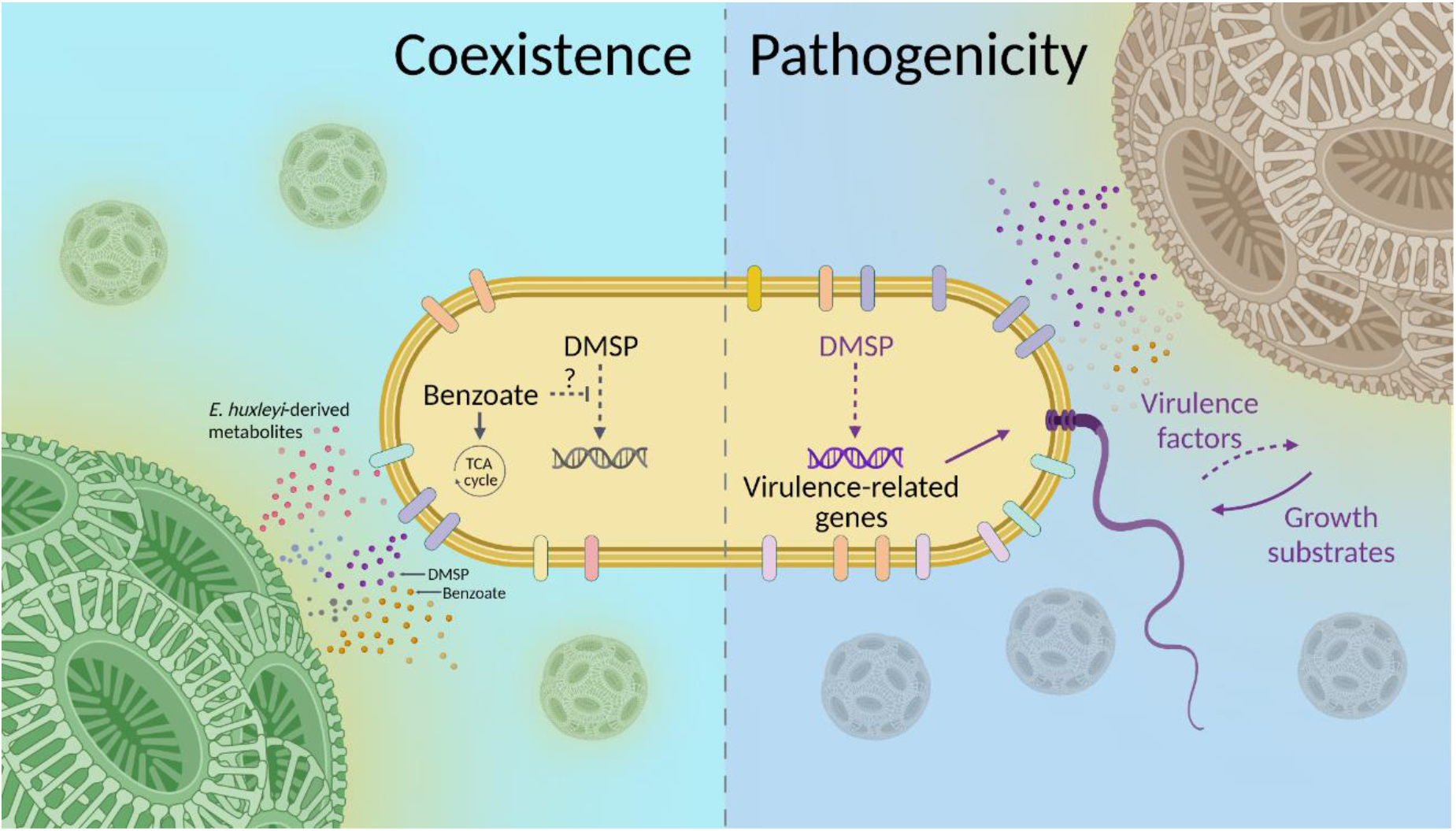
Conceptual model of the lifestyle switch of *Sulfitobacter* D7 in response to *E. huxleyi*-derived metabolites. During its interactions with *E. huxleyi, Sulfitobacter* D7 exhibits a lifestyle switch from coexistence to pathogenicity. In the coexistence phase, *E. huxleyi* secretes to the phycosphere various metabolites such as benzoate, DMSP and other growth substrates, which bacteria can uptake and consume for growth. Based on the observation that benzoate negated the pathogenicity-inducing effect of DMSP, we hypothesize that such energy-rich metabolic currencies hinder DMSP signaling in *Sulfitobacter* D7. When the algal physiological state is compromised, e.g. stationary growth, the amount of available growth substrates decreases, due to bacterial consumption and less secretion by the alga. In this context, high concentration of algal DMSP acts as a signal that alters the transcriptional profiles of the bacterium and leads to high expression of pathogenicity-related genes such as flagellar and transport genes, and yet unknown virulence factors that kill *E. huxleyi* cells. This leads to a surge of alga-derived growth substrates that are taken up efficiently by *Sulfitobacter* D7. The flagellum can mediate the dispersal of *Sulfitobacter* D7 and to forage for an alternative host.

The ability to utilize benzoate is shared among bacterial strains that are associated with *E. huxleyi* in the natural environment and in cultures (Fig. 4d) (28, 63–65). Since benzoate can act as an antibacterial compound (70, 71), we propose that secretion of benzoate by *E. huxleyi* can select for bacteria that specialize on this compound and is therefore important for the establishment of a coexistence phase. Similarly, the diatom *Asterionellopsis glacialis* produces two unique secondary metabolites, that selects for specific bacteria and also affect their behavioral response (72). Bacterial sensing of general phytoplankton-derived compounds (e.g. DMSP) together with more specific compounds (e.g. benzoate) can ensure the recognition of the algal host by the bacteria within the phycosphere. This can increase the specificity of an interaction and ensure fine-tuning of the behavior of the microorganisms by regulating gene expression. Molecular mechanisms in bacteria that integrate information perceived by various chemical signals include catabolite repression and two-component systems, which can also play a role in regulating bacterial pathogenicity (73, 74).

### The lifestyle switch of *Sulfitobacter* D7 from coexistence to pathogenicity

The experimental setup in our study demonstrated that *Sulfitobacter* D7 grown in pathogenicity-inducing media are in a different transcriptional state than in coexistence medium, which corresponds to the behavioral shift during co-culturing with *E. huxleyi* (Fig. 1). Many transport systems were differentially expressed, mainly upregulated, when *Sulfitobacter* D7 was in pathogenic state compared to the coexistence state (Fig. 3). Since bacteria often exert their pathogenicity as a mean to access nutrients released from the host, it is likely that in this mode *Sulfitobacter* D7 will maximize uptake and assimilation of metabolites released by dying *E. huxleyi* cells. High expression of transporters for branched-chain amino acids, C4 carbohydrates, DMSP, taurine and spermidine/putrescine, can facilitate the efficient uptake of these energy-rich metabolites (Fig. 3, Table S6). Upregulation of transport genes for these metabolic currencies in response to DMSP was also demonstrated in *R. pomeroyi* DSS-3, a Roseobacter often used to study bacterial metabolic exchange with phytoplankton (5, 35, 37).

During the pathogenic lifestyle there was upregulation of flagellar genes, which was functionally validated by the motility assay (Fig. 2). While DMSP is a known chemoattractant and therefore mediates the establishment of bacterial interactions with algae (33, 36), we speculate that this is not the case for *Sulfitobacter* D7 since its genome does not encode for known chemotaxis genes. We propose that the increased motility in response to DMSP in the pathogenic mode can serve as an ecological strategy to avoid competition with other bacteria in the phycosphere (75). *E. huxleyi* cell death, induced by *Sulfitobacter* D7, likely leads to a surge of intracellular metabolites that may attract other bacteria. The upregulation of flagellar motility together with transport systems can enable efficient substrate uptake by *Sulfitobacter* D7 and swimming away to forage for alternative metabolically active hosts. Such an “eat-and-run” strategy can be ecologically beneficial by facilitating the evasion from competition.

Upregulation of flagellar genes was also demonstrated during the mutualistic to pathogenic lifestyle switch of the Roseobacter *Dinoroseobacter shibae* during interactions with a dinoflagellate algal host (76). Even though *Sulfitobacter* D7 motility was increased in pathogenic mode (Fig. 2), the involvement of the flagellum may be by its alternative functions that mediate bacterial virulence (77); i.e., flagella can mediate biofilm formation and attachment to surfaces (19). Additionally, the flagellar type 3 secretion system (T3SS), which is found in the basal body and necessary for secretion of the components needed for flagellum assembly, can also be used as an export system for effector proteins in pathogenic bacteria (78). In this manner pathogenic bacteria can utilize the flagellum for multiple functions important for pathogenicity against their hosts and subsequent dispersal.

### Ecological context of bacterial lifestyle switches during algal blooms

Bacterial lifestyle switches are evident in several model systems of phytoplankton-bacteria interactions, however the ecological significance of such modes of interactions in the natural environment is elusive. In this study, we provide a contextual framework for the switch from coexistence to pathogenicity – metabolite exhaustion in the phycosphere. During a phytoplankton bloom heterotrophic bacteria can support the growth of the algae and benefit from organic matter released to the phycosphere. As the bloom progresses, various factors, such as nutrient depletion, viral infection and grazing, can compromise the algal population and its ability to provide essential metabolic currencies for optimal bacterial growth. We propose that bacteria can sense the host physiological state, by infochemicals secreted from stressed algae, and switch their behavior to pathogenic. This will result in algal cell death and bacterial proliferation, which could eventually contribute to the bloom demise. Therefore, phytoplankton-associated opportunistic bacterial pathogens constitute an underappreciated component in the regulation of algal blooms dynamics. Investigating the dynamic microscale interactions of such bacteria with phytoplankton and the metabolic crosstalk that mediate them, can provide insights into their impact on large scale biogeochemical processes in the marine environment.

## Materials and Methods

### *E. huxleyi* cultures maintenance and co-culturing with *Sulfitobacter* D7

*E. huxleyi* strains were purchased from the National Center for Marine Algae (NCMA) and maintained in filtered sea water (FSW). CCMP379, were cultured in f/2 medium (-Si) (79) and CCMP2090 was cultured in k/2 medium (-Tris, -Si) (80). Cultures were incubated at 18°C with a 16:8 h, light:dark illumination cycle. A light intensity of 100 μmol photons m^-2^ s^-1^ was provided by cool white LED lights. For all co-culturing experiments *E. huxleyi* cultures were inoculated at early exponential growth phase (4-8·10^5^ cell mL^-1^) with 10^3^ mL^-1^ *Sulfitobacter* D7 at t = 0d. When noted, DMSP or benzoate were added at t = 0d at final concentration of 100 μM.

### Enumeration of algae and bacteria abundances by flow cytometry

Flow cytometry analyses were performed on Eclipse iCyt flowcytometer (Sony Biotechnology Inc., Champaign, IL, USA) equipped with 405 and 488 nm solid-state air-cooled lasers, and with standard optic filter set-up. *E. huxleyi* cells were identified by plotting the chlorophyll fluorescence (663–737 nm) against side scatter and were quantified by counting the high-chlorophyll events. For bacterial counts, samples were fixed with a final concentration of 0.5% glutaraldehyde for at least 30 min at 4°C, then plunged into liquid nitrogen and stored at −80°C until analysis. After thawing, samples were stained with SYBR gold (Invitrogen) that was diluted 1:10,000 in Tris–EDTA buffer, incubated for 20 min at 80°C and cooled to room temperature. Samples were analyzed by flow cytometry (ex: 488 nm; em: 500–550 nm).

### Bacterial growth media

The conditioned media, Exp-CM and Stat-CM, were obtained from exponential and stationary *E. huxleyi* CCMP379 mono-cultures (Table S1), respectively, by gentle gravity filtration on GF/C filters. This method was chosen to prevent lysis of algal cells during the procedure and thus ensuring that only extracellular algae-derived metabolites, infochemicals and other components will reside in the media. CM were subsequently filtered through 0.22 μm. Exp-CM and Stat-CM were harvested on the same day of the experiment. When indicated, 100 μM DMSP was added to Exp-CM, herein Exp-CM+DMSP. MM was based on artificial sea water (ASW) (81) supplemented with basal medium (-Tris) (BM, containing essential nutrients) (82), vitamin mix (83), 0.5 mM NaNO3 and metal mix of k/2 medium (80). For the transcriptome experiment, MM were supplemented with 1 gr L^-1^ glycerol. When indicated, 100 μM DMSP was added to MM, herein MM+DMSP. For the experiment presented in Fig. 4c, 100 μM benzoate were added as sole carbon source.

### Bacterial inoculation into growth media and *E. huxleyi* cultures

Bacteria were inoculated into marine broth (Difco 2216) from a glycerol stock (kept at -80°C) and grown over-night at 28°C, 160 rpm. Bacteria were washed three times in FSW or ASW by centrifugation (10,000 g, 1 min). Bacterial inocula were counted by flow cytometry and 10^4^ bacteria mL^-1^ were inoculated to CM or MM, and 10^3^ bacteria mL^-1^ were inoculated to *E. huxleyi* cultures.

### *Sulfitobacter* D7 Transcriptome

#### Library preparation and sequencing

Experimental setup is elaborated in Fig. 1b. Samples for bacterial growth and RNA were taken at t = 24h. Bacterial cell pellets were obtained from 200 mL (MM treatments) or 160 mL (CM treatments) cultures by 2-step centrifugation: 10,000g, 10 min followed by 14,000g, 10 min, all at 4°C. Pellets were flash frozen in liquid nitrogen and stored at -80°C until further analysis. RNA extraction was carried out using the RNeasy Plant Mini Kit (Qiagen, Hilden, Germany). For disruption of cell pellets we used the OmniLyse lysis kit (Claremontbio). The rest of the RNA extraction protocol was according to manufacturer’s instruction. Library preparation was carried out according to the RNA-seq protocol developed by Avraham *et al*., (50). Briefly, DNA was removed using TURBO DNase (Ambion), RNA was fragmented and phosphorylated (at the 3’prime) using FastAP thermosensitive alkaline phosphatase (Thermo Scientific). RNA from each sample was ligated with unique RNA barcoded adaptors at the 3’, ensuring the strandedness of each transcript, using T4 RNA ligase 1 (NEB). RNA samples were pooled and treated with RiboZero (Gram-Negative Bacteria) kit (Illumina, San Diego, CA, USA) following manufacturer’s instructions in order to remove ribosomal RNA. Samples were reversed transcribed using AffinityScript RT Enzyme (Agilent) to form cDNA, and amplified by PCR. The libraries were sequenced at the Weizmann Institute of Science Core Facilities on an Illumina NextSeq500 high output v2 kit (paired end, 150 cycles).

#### Transcriptome analysis

Raw reads (64.5 million) were quality trimmed using Cutadapt (84) (-q 20 -m 20) in addition to removal of adapters. Reads were mapped to *Sulfitobacter* D7 genome assembly (GCA_003611275.1) using Bowtie2 (85) in end-to-end mode, and reads were counted on genes using HTseq, in the strict mode (86). Final reads per sample can be found in Table S12. Gene expression was quantified using DESeq2 (87) (Table S2). Differentially expressed genes were selected as genes with adjusted *P*-value <0.05, and |fold change|>2, and basemean >10 (the average of the normalized count values, dividing by size factors, taken over all samples). Priciple component analysis and similarity between samples were calculated using DESeq2 and visualized using RStudio 3.5.0. Heatmaps of gene expression were calculated using the log-normalized expression values (rld), with row standardization (scaling the means of a row to zero, with standard deviation of 1), and visualized using Partek Genomics Suite 7.0 software, Heatmapper (88) and Excel. The data has been deposited in NCBI’s Gene Expression Omnibus (GEO) and is available through GEO series accession number GSE193203.

### Functional enrichment in KEGG Pathways

Differentially expressed genes in the comparisons Exp-CM+DMSP vs. Exp-CM, Stat-CM vs. Exp-CM and MM+DMSP vs. MM were clustered using k-means analysis. For each cluster, enriched KEGG pathways (with Padj <0.01) were calculated by g:Profiler (89), using a customized reference which was constructed from *Sulfitobacter* D7 specific KEGG pathways.

### *Sulfitobacter* D7 genome mining and manual annotation

The automatic NCBI Prokaryotic Genome Annotation Pipeline was used for *Sulfitobacter* D7 genome functions prediction (43). We manually validated the function of genes related to DMSP metabolism, transport, benzoate degradation and flagella assembly by cross examining their annotation using KEGG, COG and IMG/M. For genes with no or inconsistent annotation, we also searched for functional domains using the Conserved Domain Database (CDD), and we ran BLAST using genes with known functions to validate the annotation.

#### Transport genes

The automatic annotations of transport genes were manually validated by ensuring that the genes were annotated as such by at least two automatic annotation platforms and by CDD search. Since transport systems are organized in operon-like structures, we examined the genes adjacent to the transport genes and manually annotated these additional transport genes. The transporters presented in the heatmap (Fig. 3) are the full transport systems that at least two of the genes in each system were DE. The Venn diagrams (Fig. S2) contains only the transport genes that were significantly DE. The substrates for each transport system were inferred automatically, therefore, the exact substrates were not experimentally validated.

#### Benzoate degradation genes

*Sulfitobacter* D7 benzoate degradation pathway was reconstructed using the KEGG mapping tool (90). All the genes in the catechol branch of benzoate degradation (Fig. S3) were found, except Muconolactone isomerase *CatC*. For the visualization of the organization of the genetic locus of benzoate-related genes we utilized the IMG/M platform (91).

#### Flagellar genes

We manually validated the annotation of all the flagella genes in *Sulfitobacter* D7 genome and found most of the genes, except for three: *FliQ, FliJ* and *FliD* (92). For the visualization of the organization of the genetic locus of flagellar genes we utilized the IMG/M platform (91).

### Bacterial motility assay

Motility was assessed by examining the expansion of bacterial colonies plated on semi-solid agar (93). Semi-solid media of Exp-CM, Exp-CM+DMSP and Stat-CM were prepared by mixing boiling sterile 3% agarose with CM, which was pre-heated to ∼50°C, in a 1:9 ratio (final concentration of 0.3% agarose). Media was quickly distributed in 6-well plates, ∼5 mL per plate, and was left to solidify for ∼ 1 h. *Sulfitobacter* D7 were pre-grown in liquid Exp-CM, Exp-CM+DMSP and Stat-CM in order to induce the appropriate expression of flagellar genes. For control, bacteria were pre-grown in liquid ½MB, lacking algal DMSP and infochemicals. After 24 h bacterial abundance was evaluated and the concentration of bacteria in each media was normalized to 2·10^6^ mL^-1^, to ensure that the difference in colony size would be indicative of motility and not abundance of bacteria. Bacteria grown in CM were plated on the corresponding semi-solid CM (0.3% agarose) and bacteria grown in ½MB were plated on each semi-solid CM. For plating, 1 μL of bacteria were pipetted in the center of each well containing semi-solid media, in 10-12 replicates per treatment. Colonies were visualized with 2X magnification after 6 days using Nikon SMZ18 Steriomicroscope. Colonies measurements were performed using the Annotation and Measurements tool of the Nikon NIS-Elements Analysis D software.

### Quantification of benzoate in media extracts

To quantify extracellular benzoate concentrations, cultures were filtered gently over 0.22 μm filters, acidified, and led through hydrophilic-lipophilic balanced solid phase extraction (SPE) cartridges, as described in Kuhlisch et al. (94). Glassware and chemically resistant equipment were used whenever possible and cleaned with HCl (1% or 10%) and Deconex 20 NS-x (Borer Chemie, Zuchwil, Switzerland) to reduce contaminations. Per sample, 50 mL (bacterial cultures) or 300 mL (algal cultures) of filtrate was collected in glass Erlenmeyer flasks and spiked with 5 μL of benzoate-d_5_ (98%, Cambridge Isotope Laboratries, Tewsbury, MA, USA; 1.276 μg/μL in MeOH) for 1 μM final concentration, as internal standard. The filtrates were incubated for 30 min and then acidified to pH 2.0 using 10% HCl. Metabolites were extracted using hydrophilic-lipophilic balanced SPE cartridges (Oasis HLB, 200 mg, Waters, Milford, MA, USA) as follows: cartridges were conditioned (6 mL methanol), equilibrated (6 mL 0.01 N HCl), and then loaded by gravity with the acidified samples (45 min). The cartridges were then washed (18 mL 0.01 N HCl), dried completely using a vacuum pump, and gravity-eluted with 2× 2 mL methanol into 4 mL glass vials. Eluates were stored at -20°C, dried under a flow of nitrogen at 0.5 mL/min and 30°C (TurboVap LV, Biotage, Uppsala, Sweden), and stored at -20°C until further processing. Dried extracts were thawed, re-dissolved in 300 μL methanol:water (1:1, v:v), vortexed, sonicated for 10 min, and centrifuged at 3,200×g for 10 min at 4°C. The supernatants were transferred to 200 μL glass inserts in autosampler vials and analyzed by ultra-high-performance liquid chromatography-electrospray-high resolution mass spectrometry (UHPLC-ESI-HRMS). An aliquot of 1 μL was analyzed using UPLC coupled to a photodiode detector (ACQUITY UPLC I-Class, Waters) and a quadrupole time-of-flight (QToF) mass spectrometer (SYNAPT G2 HDMS, Waters), as described previously with slight modifications. Chromatographic separation was carried out using an ACQUITY UPLC BEH C18 column (100 × 2.1 mm, 1.7 μm; Waters) attached to a VanGuard pre-column (5 × 2.1 mm, 1.7 μm; Waters). The mobile phase, at a flow rate of 0.3 mL/min, consisted of water (mobile phase A) and acetonitrile (mobile phase B), both with 0.1% formic acid, and set as follows: a liner gradient from 100-75% A in 20 min, from 75-0% A in 6 min, 2 min of 100% B, and 2 min to return to the initial conditions and re-equilibrate the column. The PDA detector was set to 200-600 nm. A divert valve (Rheodyne) excluded 0-1 min and 25.5-30 min from injection to the mass spectrometer. The ESI source was operated in negative ionization mode and set to 140°C source and 450°C desolvation temperature, 1.0 kV capillary voltage, and 27 eV cone voltage, using nitrogen as desolvation gas (800 L/h) and cone gas (10 L/h). The mass spectrometer was operated in full scan MSE resolution mode with a mass range of 50-1600 Da and the mass resolution tuned to 23,000 at m/z 554 alternating with 0.1 min scan time between low-(4 eV collision energy) and high-energy scan function (collision energy ramp of 15-50 eV).

An external calibration curve was processed in parallel. Aliquots of 100 mL artificial seawater (ASW) were spiked with 10 μL of d5-benzoate as internal standard (IS) (1 μM final concentration) and benzoate standard solutions to reach final concentrations of 0.2, 1, 2, 10, 20, and 100 μM benzoate. Two blanks were prepared, one blank that was spiked only with the IS, and one blank lacking both IS and benzoate. Each sample was divided to duplicates of 50 mL and extracted as described above. After re-dissolving in 200 μL methanol:water (1:1, v:v), samples were injected subsequent to the biological samples. The peak areas of the [M-H]-ions for the IS (*m*/*z* 126.06) and benzoate (*m*/*z* 121.029) were extracted above a signal-to-noise threshold of 10 using QuanLynx (Version 4.1, Waters), and the analyte response calculated by dividing the area of benzoate by the IS (Fig. S5). The response (y) was then plotted against the concentration of benzoate (x), and the slope and intercept for a linear regression calculated (y = 0.4741x + 1.665, R^2^ = 0.99) (Fig. S5). Quantification of the samples was done based on the analyte response in each sample and the calibration curve. The limit of quantification was 200 nM.

### Prevalence of benzoate transport and catabolism genes in genomes of phytoplankton-associated bacteria

Bacterial benzoate degradation pathways and the genes encoding for the metabolic enzymes were reconstructed with the use of MetaCyc (95) and the KEGG Pathway database (96) (Fig. S3). Selected genes, encoding for benzoate transporters and for the enzymes mediating the initial steps of benzoate metabolism in each pathway, were used to search for similar proteins in bacterial genomes using BLASTp. The list of these query genes, which were all previously experimentally validated, is found in Table S10. The target bacterial genomes were selected based on their known association with *E. huxleyi* and other phytoplankton species (Table S9). Positive hits had an E-value <0.005, identity > 30% and coverage >30. Hits with lower coverage and/or identity were considered as “Partial”. The results are summarized in Table S9.

### Statistical analyses

For the motility assay (Fig. 2b) we used 2-way ANOVA, followed by Tukey’s post-hoc test, using the R-package “emmeans”. For benzoate consumption (Fig. 4c) we used a mixed effects model, with treatment and time as fixed effects, and replicate as a random effect, using the R package’s “lme4” and “lmerTest”. For the *E. huxleyi* growth curves (Fig. 5 a,b) we used a mixed effects model, with treatment, bacteria (none or D7) and time as fixed effects, and replicate as a random effect. For the bacterial growth curves (Fig. 5c) we used a mixed effects model, with treatment and time as fixed effects, and replicate as a random effect. Slopes within the mixed models were compared using the ‘emmeans’ package. *P*-values for all comparisons are presented in Table S11. All analyses were done using R, v. 4.1.2.

### Figures preparation

Figures and illustrations were prepared using PowerPoint, Excel and <u>BioRender.com</u>.

## Acknowledgments

We thank Ron Rotkopf for his assistance in statistical analysis. We thank Daniella Schatz for constructive feedback and scientific discussions. We thank Assaf R. Gavish for fruitful discussions and assistance in graphics. This research was supported by the European Research Council CoG (VIROCELLSPHERE grant no. 681715) and research grants from the Estate of Bernard Berkowitz and the *de Botton* Center for Marine Science awarded to Assaf Vardi.

## References

1. C. B. Field, M. J. Behrenfeld, J. T. Randerson, P. Falkowski, Primary Production of the Biosphere: Integrating Terrestrial and Oceanic Components. Science 281, 237–240 (1998).

2. J. R. Seymour, S. A. Amin, J.-B. Raina, R. Stocker, Zooming in on the phycosphere: the ecological interface for phytoplankton–bacteria relationships. Nat. Microbiol. 2, 17065 (2017).

3. E. Cirri, G. Pohnert, Algae−bacteria interactions that balance the planktonic microbiome. New Phytol. 223, 100–106 (2019).

4. W. Bell, R. Mitchell, Chemotactic and growth responses of marine bacteria to algal extracellular products. Biol. Bull. 143, 265–277 (1972).

5. M. Landa, A. S. Burns, S. J. Roth, M. A. Moran, Bacterial transcriptome remodeling during sequential co-culture with a marine dinoflagellate and diatom. ISME J. 11, 2677–2690 (2017).

6. E. Segev, et al., Dynamic metabolic exchange governs a marine algal-bacterial interaction. Elife 5, e17473 (2016).

7. H. Wang, J. Tomasch, M. Jarek, I. Wagner-Döbler, A dual-species co-cultivation system to study the interactions between Roseobacters and dinoflagellates. Front. Microbiol. 5, 311 (2014).

8. S. A. Amin, et al., Photolysis of iron-siderophore chelates promotes bacterial-algal mutualism. Proc. Natl. Acad. Sci. 106, 17071–17076 (2009).

9. M. T. Croft, A. D. Lawrence, E. Raux-Deery, M. J. Warren, A. G. Smith, Algae acquire vitamin B<sub>12 </sub>through a symbiotic relationship with bacteria. Nature 438, 90–93 (2005).

10. G. Pohnert, M. Steinke, R. Tollrian, Chemical cues, defence metabolites and the shaping of pelagic interspecific interactions. Trends Ecol. Evol. 22, 198–204 (2007).

11. M. R. Seyedsayamdost, R. J. Case, R. Kolter, J. Clardy, The Jekyll-and-Hyde chemistry of Phaeobacter gallaeciensis. Nat. Chem. 3, 331–335 (2011).

12. S. A. Amin, et al., Interaction and signalling between a cosmopolitan phytoplankton and associated bacteria. Nature 522, 98–101 (2015).

13. N. Barak-Gavish, et al., Bacterial virulence against an oceanic bloom-forming phytoplankter is mediated by algal DMSP. Sci. Adv. 4, eaau5716 (2018).

14. R. N. Slightom, A. Buchan, Surface Colonization by Marine Roseobacters: Integrating Genotype and Phenotype. Appl. Environ. Microbiol. 75, 6027–6037 (2009).

15. R. Stocker, J. R. Seymour, Ecology and physics of bacterial chemotaxis in the ocean. Microbiol. Mol. Biol. Rev. 76, 792–812 (2012).

16. G. Furusawa, T. Yoshikawa, A. Yasuda, T. Sakata, Algicidal activity and gliding motility of Saprospira sp. SS98-5. Can. J. Microbiol. 49, 92–100 (2003).

17. T. R. Miller, R. Belas, Motility is involved in Silicibacter sp. TM1040 interaction with dinoflagellates. Environ. Microbiol. 8, 1648–1659 (2006).

18. E. C. Sonnenschein, D. A. Syit, H.-P. Grossart, M. S. Ullrich, Chemotaxis of Marinobacter adhaerens and its impact on attachment to the diatom Thalassiosira weissflogii. Appl. Environ. Microbiol. 78, 6900–7 (2012).

19. Y. Li, et al., Chitinase producing bacteria with direct algicidal activity on marine diatoms. 1–13 (2016).

20. C. Fei, et al., Quorum sensing regulates ‘swim-or-stick’ lifestyle in the phycosphere. Environ. Microbiol. 22, 4761–4778 (2020).

21. X. Mayali, P. J. S. Franks, F. Azam, Cultivation and ecosystem role of a marine Roseobacter clade-affiliated cluster bacterium. Appl. Environ. Microbiol. 74, 2595–2603 (2008).

22. A. Buchan, G. R. LeCleir, C. A. Gulvik, J. M. González, Master recyclers: features and functions of bacteria associated with phytoplankton blooms. Nat. Rev. Microbiol. 12, 686–698 (2014).

23. B. Rink, et al., Effects of phytoplankton bloom in a coastal ecosystem on the composition of bacterial communities. Aquat. Microb. Ecol. 48, 47–60 (2007).

24. J. M. González, M. A. Moran, Numerical dominance of a group of marine bacteria in the alpha-subclass of the class Proteobacteria in coastal seawater. Appl. Environ. Microbiol. 63, 4237–4242 (1997).

25. M. Alavi, T. Miller, K. Erlandson, R. Schneider, R. Belas, Bacterial community associated with Pfiesteria-like dinoflagellate cultures. Environ. Microbiol. 3, 380–396 (2001).

26. S. A. Amin, M. S. Parker, E. V. Armbrust, Interactions between diatoms and bacteria. Microbiol. Mol. Biol. Rev. 76, 667–684 (2012).

27. G. Behringer, et al., Bacterial communities of diatoms display strong conservation across strains and time. Front. Microbiol. 9, 1–15 (2018).

28. F. Vincent, et al., Viral infection switches the balance between bacterial and eukaryotic recyclers of organic matter during algal blooms. bioRxiv, 2021.10.25.465659 (2021).

29. H. Geng, R. Belas, Molecular mechanisms underlying roseobacter-phytoplankton symbioses. Curr. Opin. Biotechnol. 21, 332–8 (2010).

30. R. J. Newton, et al., Genome characteristics of a generalist marine bacterial lineage. ISME J. 4, 784–798 (2010).

31. M. D. Keller, Dimethyl Sulfide Production and Marine Phytoplankton: The Importance of Species Composition and Cell Size. Biol. Oceanogr. 6, 375–382 (1989).

32. T. R. Miller, R. Belas, Dimethylsulfoniopropionate metabolism by Pfiesteria-associated Roseobacter spp. Appl. Environ. Microbiol. 70, 3383–3391 (2004).

33. T. R. Miller, K. Hnilicka, A. Dziedzic, P. Desplats, R. Belas, Chemotaxis of Silicibacter sp. strain TM1040 toward dinoflagellate products. Appl. Environ. Microbiol. 70, 4692–4701 (2004).

34. P. Sule, R. Belas, A novel inducer of Roseobacter motility is also a disruptor of algal symbiosis. J. Bacteriol. 195, 637–46 (2013).

35. H. Bürgmann, et al., Transcriptional response of Silicibacter pomeroyi DSS-3 to dimethylsulfoniopropionate (DMSP). Environ. Microbiol. 9, 2742–2755 (2007).

36. J. R. Seymour, R. Simó, T. Ahmed, R. Stocker, Chemoattraction to dimethylsulfoniopropionate throughout the marine microbial food web. Science 329, 342–345 (2010).

37. B. P. Durham, et al., Cryptic carbon and sulfur cycling between surface ocean plankton. Proc. Natl. Acad. Sci. 112, 453–457 (2015).

38. M. B. Cooper, et al., Cross-exchange of B-vitamins underpins a mutualistic interaction between Ostreococcus tauri and Dinoroseobacter shibae. ISME J. 13, 334–345 (2019).

39. A. R. Bramucci, et al., The Bacterial Symbiont Phaeobacter inhibens Shapes the Life History of Its Algal Host Emiliania huxleyi. Front. Mar. Sci. 5, 188 (2018).

40. T. J. Mayers, A. R. Bramucci, K. M. Yakimovich, R. J. Case, A Bacterial Pathogen Displaying Temperature-Enhanced Virulence of the Microalga Emiliania huxleyi. Front. Microbiol. 7, 892 (2016).

41. C. J. S. Bolch, T. A. Bejoy, D. H. Green, Bacterial Associates Modify Growth Dynamics of the Dinoflagellate Gymnodinium catenatum. Front. Microbiol. 8, 670 (2017).

42. A. Shemi, et al., Dimethyl sulfide mediates microbial predator–prey interactions between zooplankton and algae in the ocean. Nat. Microbiol. 6, 1357–1366 (2021).

43. C. Ku, N. Barak-Gavish, M. Maienschein-Cline, S. J. Green, A. Vardi, Complete Genome Sequence of Sulfitobacter sp. Strain D7, a Virulent Bacterium Isolated from an Emiliania huxleyi Algal Bloom in the North Atlantic. Microbiol. Resour. Announc. 7, e01379–18 (2018).

44. N. Y. D. Ankrah, T. Lane, C. R. Budinoff, M. K. Hadden, A. Buchan, Draft Genome sequence of Sulfitobacter sp. CB2047, a member of the Roseobacter clade of marine bacteria, isolated from an Emiliania huxleyi Bloom. Genome Announc. 2, e01125–14 (2014).

45. E. C. Howard, et al., Bacterial taxa that limit sulfur flux from the ocean. Science 314, 649–652 (2006).

46. J. D. Todd, et al., Structural and regulatory genes required to make the gas dimethyl sulfide in bacteria. Science 315, 666–9 (2007).

47. C. R. Reisch, et al., Novel pathway for assimilation of dimethylsulphoniopropionate widespread in marine bacteria. Nature 473, 208–211 (2011).

48. L. Sun, A. R. J. Curson, J. D. Todd, A. W. B. Johnston, Diversity of DMSP transport in marine bacteria, revealed by genetic analyses. Biogeochemistry 110, 121–130 (2012).

49. A. R. J. Curson, J. D. Todd, M. J. Sullivan, A. W. B. Johnston, Catabolism of dimethylsulphoniopropionate: microorganisms, enzymes and genes. Nat. Rev. Microbiol. 9, 849–859 (2011).

50. R. Avraham, et al., A highly multiplexed and sensitive RNA-seq protocol for simultaneous analysis of host and pathogen transcriptomes. Nat. Protoc. 11, 1477–1491 (2016).

51. S. D. Cook, An Historical Review of Phenylacetic Acid. Plant Cell Physiol. 60, 243–254 (2019).

52. V. Thiel, et al., Identification and biosynthesis of tropone derivatives and sulfur volatiles produced by bacteria of the marine Roseobacter clade. Org. Biomol. Chem. 8, 234–246 (2010).

53. R. Wang, É. Gallant, M. R. Seyedsayamdost, Investigation of the genetics and biochemistry of roseobacticide production in the Roseobacter clade bacterium Phaeobacter inhibens. MBio 7, e02118–15 (2016).

54. F. X. Ferrer-González, et al., Resource partitioning of phytoplankton metabolites that support bacterial heterotrophy. ISME J. 15, 762–773 (2021).

55. T. Obata, et al., Gas-Chromatography Mass-Spectrometry (GC-MS) Based Metabolite Profiling Reveals Mannitol as a Major Storage Carbohydrate in the Coccolithophorid Alga Emiliania huxleyi. Metabolites 3, 168–184 (2013).

56. Y. Tsuji, I. Suzuki, Y. Shiraiwa, Enzymological Evidence for the Function of a Plastid-Located Pyruvate Carboxylase in the Haptophyte alga Emiliania huxleyi: A Novel Pathway for the Production of C4 Compounds. Plant Cell Physiol. 53, 1043–1052 (2012).

57. A. Buchan, E. L. Neidle, M. A. Moran, Diverse Organization of Genes of the β-Ketoadipate Pathway in Members of the Marine Roseobacter Lineage. Appl. Environ. Microbiol. 70, 1658–1668 (2004).

58. M. A. Moran, et al., Ecological genomics of marine roseobacters. Appl. Environ. Microbiol. 73, 4559–4569 (2007).

59. C. S. Harwood, R. E. Parales, The β-ketoadipate pathway and the biology of self-identity. Annu. Rev. Microbiol. 50, 553–590 (1996).

60. G. Fuchs, M. Boll, J. Heider, Microbial degradation of aromatic compounds — from one strategy to four. Nat. Rev. Microbiol. 9, 803–816 (2011).

61. L. S. Collier, G. L. Gaines, E. L. Neidle, Regulation of Benzoate Degradation in Acinetobacter sp. Strain ADP1 by BenM, a LysR-Type Transcriptional Activator. J. Bacteriol. 180, 2493–2501 (1998).

62. B. M. Bundy, L. S. Collier, T. R. Hoover, E. L. Neidle, Synergistic transcriptional activation by one regulatory protein in response to two metabolites. Proc. Natl. Acad. Sci. 99, 7693–7698 (2002).

63. A. R. R. Rosana, et al., Draft Genome Sequences of Seven Bacterial Strains Isolated from a Polymicrobial Culture of Coccolith-Bearing (C-Type) Emiliania huxleyi M217. Genome Announc. 4, 9–10 (2016).

64. D. H. Green, V. Echavarri-bravo, D. Brennan, M. C. Hart, Bacterial diversity associated with the coccolithophorid algae Emiliania huxleyi and Coccolithus pelagicus f. braarudii. Biomed Res. Int. 2015 (2015).

65. F. D. Orata, et al., Draft Genome Sequences of Four Bacterial Strains Isolated from a Polymicrobial Culture of Naked (N-Type) Emiliania huxleyi CCMP1516. Genome Announc. 4, 9–10 (2016).

66. A. R. J. Curson, et al., Dimethylsulfoniopropionate biosynthesis in marine bacteria and identification of the key gene in this process. Nat. Microbiol. 2, 17009 (2017).

67. A. Jensen, Excretion of organic carbon as function of nutrient stress. Mar. Phytoplankt. Product. Proc. Symp. Taormina, 1983, 61–72 (1984).

68. A. Barofsky, C. Vidoudez, G. Pohnert, Metabolic profiling reveals growth stage variability in diatom exudates. Limnol. Oceanogr. Methods 7, 382–390 (2009).

69. I. Obernosterer, G. J. Herndl, Phytoplankton extracellular release and bacterial growth: Dependence on the inorganic N:P ratio. Mar. Ecol. Prog. Ser. 116, 247–258 (1995).

70. H. Haque, T. J. Cutright, B.-M. Z. Newby, Effectiveness of sodium benzoate as a freshwater low toxicity antifoulant when dispersed in solution and entrapped in silicone coatings. Biofouling 21, 109–119 (2005).

71. D. H. Amin, A. Abolmaaty, Efficacy assessment of various natural and organic antimicrobials against Escherichia coli O157:H7, Salmonella enteritidis and Listeria monocytogenes. Bull. Natl. Res. Cent. 44, 172 (2020).

72. A. A. Shibl, et al., Diatom modulation of select bacteria through use of two unique secondary metabolites. Proc. Natl. Acad. Sci. 117, 27445–27455 (2020).

73. B. Görke, J. Stülke, Carbon catabolite repression in bacteria: many ways to make the most out of nutrients. Nat. Rev. Microbiol. 6, 613–624 (2008).

74. D. Beier, R. Gross, Regulation of bacterial virulence by two-component systems. Curr. Opin. Microbiol. 9, 143–152 (2006).

75. Y. Yawata, et al., Competition-dispersal tradeoff ecologically differentiates recently speciated marine bacterioplankton populations. Proc. Natl. Acad. Sci., 1–6 (2014).

76. H. Wang, et al., Identification of Genetic Modules Mediating the Jekyll and Hyde Interaction of Dinoroseobacter shibae with the Dinoflagellate Prorocentrum minimum. Front. Microbiol. 6, 1–8 (2015).

77. B. Chaban, H. V. Hughes, M. Beeby, The flagellum in bacterial pathogens: For motility and a whole lot more. Semin. Cell Dev. Biol. 46, 91–103 (2015).

78. A. Diepold, J. P. Armitage, Type III secretion systems: the bacterial flagellum and the injectisome. Philos. Trans. R. Soc. B Biol. Sci. 370, 20150020 (2015).

79. R. R. L. Guillard, J. H. Ryther, Studies of marine planktonic diatoms. I. Cyclotella nana Hustedt, and Detonula confervacea (cleve) Gran. Can. J. Microbiol. 8, 229–239 (1962).

80. M. D. Keller, R. C. Seluin, W. Claus, R. R. L. Guillard, Media for the culture of oceanic ultraphytoplankton. J. Phycol. 23, 633–638 (1987).

81. C. Goyet, A. Poisson, New determination of carbonic acid dissociation constants in seawater as a function of temperature and salinity. Deep Sea Res. Part A, Oceanogr. Res. Pap. 36, 1635–1654 (1989).

82. P. Baumann, L. Baumann, “The marine Gram-negative eubacteria: genera Photobacterium, Beneckea, Alteromonas, Pseudomonas, and Alcaligenes” in The Prokaryotes, M. P. Starr, H. Stolp, H. G. Triiper, A. Balows, H. G. Schlegel, Eds. (Springer-Verlag, 1981), pp. 1302–1331.

83. J. M. González, F. Mayer, M. A. Moran, R. E. Hodson, W. B. Whitman, Microbulbifer hydrolyticus gen. nov., sp. nov., and Marinobacterium georgiense gen. nov., sp. nov., two marine bacteria from a lignin-rich pulp mill waste enrichment community. Int. J. Syst. Bacteriol. 47, 369–376 (1997).

84. M. Martin, Cutadapt removes adapter sequences from high-throughput sequencing reads. EMBnet.journal 17, 10 (2011).

85. B. Langmead, S. L. Salzberg, Fast gapped-read alignment with Bowtie 2. Nat. Methods 9, 357–359 (2012).

86. S. Anders, P. T. Pyl, W. Huber, HTSeq — a Python framework to work with high-throughput sequencing data. Bioinformatics 31, 166–169 (2015).

87. M. I. Love, W. Huber, S. Anders, Moderated estimation of fold change and dispersion for RNA-seq data with DESeq2. Genome Biol. 15, 550 (2014).

88. S. Babicki, et al., Heatmapper: web-enabled heat mapping for all. Nucleic Acids Res. 44, W147–W153 (2016).

89. U. Raudvere, et al., g:Profiler: a web server for functional enrichment analysis and conversions of gene lists (2019 update). Nucleic Acids Res. 47, W191–W198 (2019).

90. M. Kanehisa, Y. Sato, M. Kawashima, KEGG mapping tools for uncovering hidden features in biological data. Protein Sci., 1–7 (2021).

91. I.-M. A. Chen, et al., IMG/M: integrated genome and metagenome comparative data analysis system. Nucleic Acids Res. 45, D507–D516 (2017).

92. F. F. V Chevance, K. T. Hughes, Coordinating assembly of a bacterial macromolecular machine. Nat. Rev. Microbiol. 6, 455–465 (2008).

93. A. J. Wolfe, H. C. Berg, Migration of bacteria in semisolid agar. Proc. Natl. Acad. Sci. 86, 6973–6977 (1989).

94. C. Kuhlisch, et al., Viral infection of algal blooms leaves a unique metabolic footprint on the dissolved organic matter in the ocean. Sci. Adv. 7, 1–14 (2021).

95. R. Caspi, et al., The MetaCyc database of metabolic pathways and enzymes and the BioCyc collection of Pathway/Genome Databases. Nucleic Acids Res. 42, D459–D471 (2014).

96. M. Kanehisa, KEGG: Kyoto Encyclopedia of Genes and Genomes. Nucleic Acids Res. 28, 27–30 (2000).

