## Supplementary Information for "Bacterial lifestyle switch in response to algal metabolites"

**This PDF file includes:**

SI Figures S1 to S5
SI Tables S1 to S12 (Table S2 and S6 are in separate files)
SI References

### SI Figures

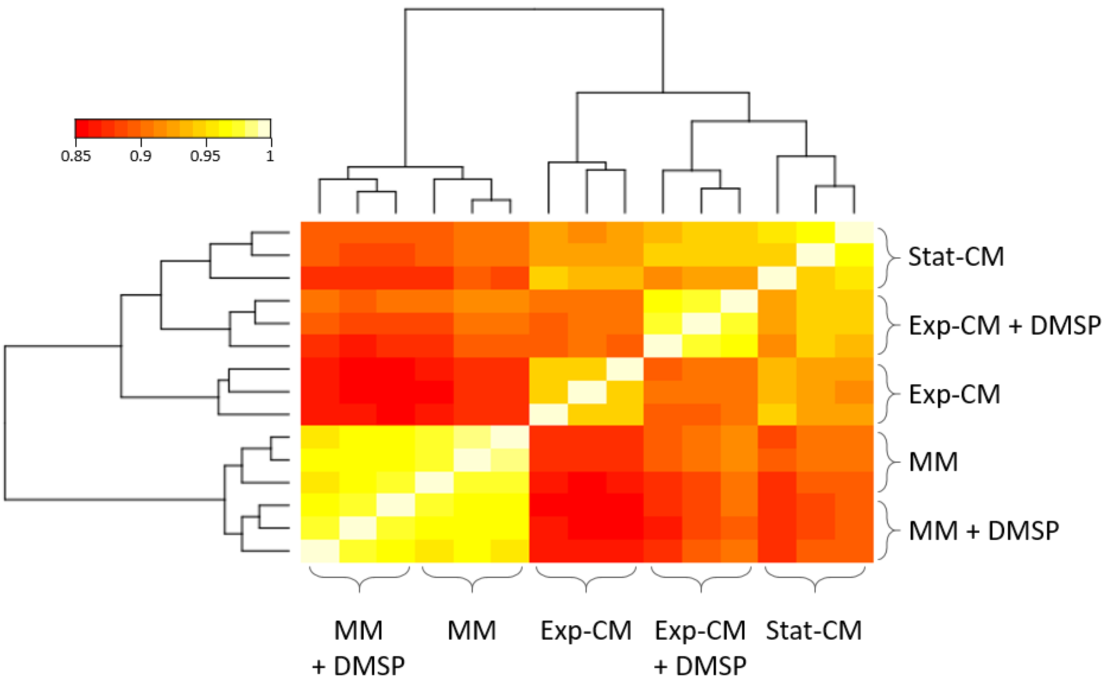


**<u>Figure S1.</u> Gene expression similarity between the treatments of *Sulfitobacter* D7 transcriptome.**

Pearson correlation and dendogram of hierarchical clustering between all samples in *Sulfitobacter* D7 transcriptome according to gene expression values. Replicates of same treatment show high correlation and cluster together. Conditioned media (CM) and minimal media (MM) treatments cluster separately.

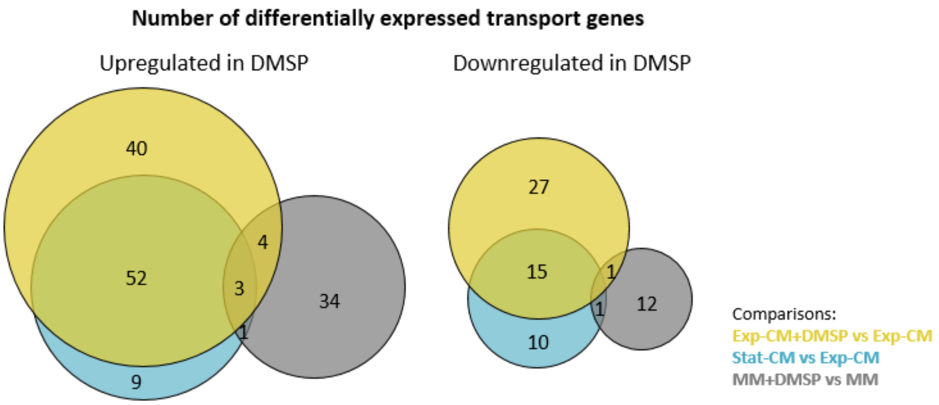


**Figure S2. Differential expression of *Sulfitobacter* D7 transport genes in coexistence and pathogenic states and in response to DMSP.**

Venn diagram showing the number of differentially expressed transport genes in the indicated treatment comparisons. The left diagram corresponds to genes that are upregulated in samples with higher DMSP concentration, and the right side are downregulated genes. Venn diagrams were generated using BioVenn (1).

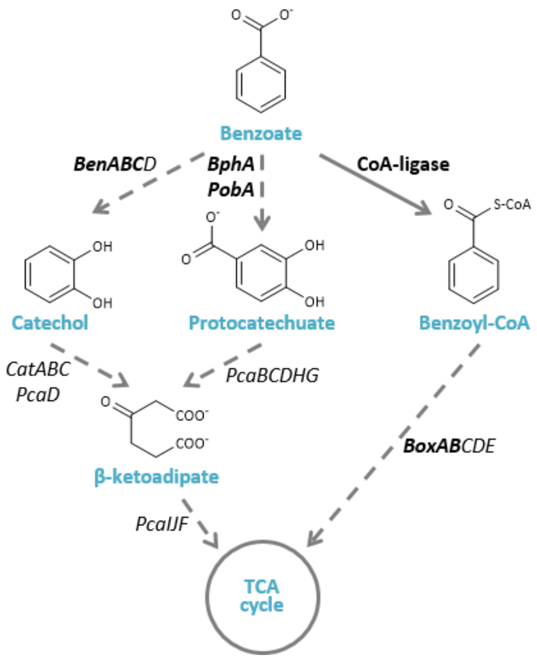


**Figure S3. Bacterial benzoate degradation pathways.**
Aerobic degradation of benzoate by bacteria can occur through three possible pathways: through catechol (*Ben* genes), through protocatecuate (*BphA and PobA* genes) and through benzoyl-CoA (*Box* genes). The first two pathways converge to β-ketoadipate, and in all the three pathways the end products are acetyl-CoA and succinate/succinyl-CoA, which are channeled to the TCA cycle (2). The names of the genes encoding for the enzymes mediating the metabolic transformations are denoted next to the arrows. Full arrow represent a single enzymatic reaction, while dashed arrows represent multiple enzymatic reactions. The genes marked in bold were used as query genes for examining the prevalence of benzoate transport and catabolism genes in genomes of phytoplankton-associated bacteria (Fig. 4d, Table S9 and S10).

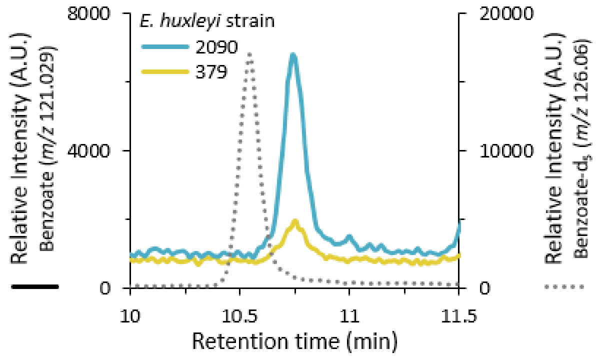


**Figure S4.** **Detection of benzoate in medium extracts of *E. huxleyi* cultures.**
Extracted ion chromatograms (EICs) of the molecular ions $[M\text{-}H]^-$ of benzoate (*m*/*z* 121.029) and the internal standard (IS) benzoate-$d_5$ (*m*/*z* 126.06) in solid phase extracts of cell-free filtrates derived from cultures of *E. huxleyi* strains CCMP2090 (late exponential phase) and CCMP379 (early stationary phase), spiked with IS at 1 µM. The EIC of the IS depicts the relative intensity in the CCMP2090 culture sample. The difference in retention times compared to Fig. S5 is due to slight modifications in the chromatographic setup (see Materials and Methods).

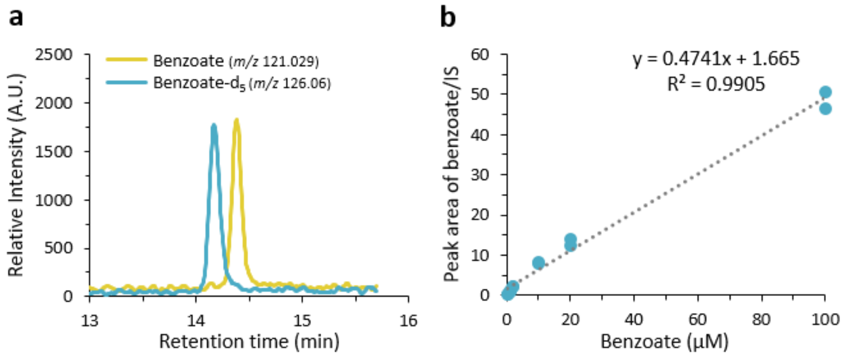


__Figure S5.__ **Detection and quantification of benzoate in artificial seawater extracts by ultra-high-performance liquid chromatography-electrospray-high resolution mass spectrometry (UHPLC-ESI-HRM).**

(a) Extracted ion chromatograms of the molecular ions $[M\text{-}H]^-$ of benzoate (*m*/*z* 121.029) and benzoate-$d_5$ (*m*/*z* 126.06) as internal standard from a solid phase extract of artificial seawater that was spiked with both metabolites at 1 µM. The internal standard benzoate-$d_5$ elutes slightly earlier due to complete deuterium labeling of the aromatic ring. (b) Standard calibration curve of sodium benzoate and benzoate-$d_5$ as internal standard (IS) from solid phase extracts of artificial seawater samples that were spiked in duplicates at different benzoate concentrations ranging from 0.2 µM-100 µM.

### SI Tables

**Table S1. Conditions of the media used for *Sulfitobacter* D7 transcriptome.**

| Media | *Sulfitobacter* D7 abundance at t=24h ($10^6 \cdot mL^{-1}$)[b] | DMSP concentration (µM)[c] | For conditioned media (CM)[a] Days of *E. huxleyi* growth | *E. huxleyi* abundance ($cells \cdot mL^{-1}$) | % Dead cells[c] |
|---|---|---|---|---|---|
| Exp-CM | 2.6 ± 0.1 | 3 | 5 | $1 \cdot 10^6$ | 8.5 % |
| Exp-CM+DMSP | 3.7 ± 0.3 | 100 | 5 | $1 \cdot 10^6$ | 8.5 % |
| Stat-CM | 17.5 ± 0.8 | 10.5 | 12 | $3 \cdot 10^6$ | 18 % |
| MM | 7.5 ± 0.5 | 0 | – | – | – |
| MM+DMSP | 18.6 ± 1.3 | 100 | – | – | – |

[a] Conditions of exponential and stationary *E. huxleyi* cultures from which Exp-CM and Stat-CM were obtained, respectively (n=1)

[b] Results represent average ± SD (n=3). At t=24 h bacteria were harvested for transcriptome analysis (experimental setup, Fig. 1b)

[c] DMSP and cell death measurements are described in Barak-Gavish *et al.,* 2018 (n=1)

**Table S2.** **RNA sequencing data of *Sulfitobacter* D7 transcriptome.**

File attached.

**<u>Table S3.</u> Number of *Sulfitobacter* D7 differentially expressed (DE) genes in pathogenicity vs. coexistence modes and in response to DMSP.**

| | Exp-CM+DMSP vs. Exp-CM | Stat-CM vs. Exp-CM | MM+DMSP vs. MM |
|---|---|---|---|
| Upregulated | 560 | 358 | 107 |
| Downregulated | 408 | 237 | 63 |
| Total | 968 | 495 | 170 |

DE genes were defined as genes with |fold change|>2 and adjusted *P*-value ≤ 0.05

**Table S4.** ***Sulfitobacter*** **D7 expression of genes belonging to KEGG Pathways that are enriched in clusters 2-4 of the heatmap in Fig. 1d.**

| Cluster[a] | KEGG Pathway enriched | Locus tag | Annotation[b] | Normalized expression values (log transformed) (triplicate samples for each treatment) | | | | | | | | | | | | | | | Differential expression[c] Fold change ($log_2$) | | |
|---|---|---|---|---|---|---|---|---|---|---|---|---|---|---|---|---|---|---|---|---|---|
| | | | | Exp-CM | | | Exp-CM +DMSP | | | Stat-CM | | | MM | | | MM +DMSP | | | Exp-CM +DMSP vs. Exp-CM | Stat-CM vs. Exp-CM | MM +DMSP vs. MM |
| 2 | Phenyl-alanine metabolism | B5M07_01040 | Phenylacetic acid degradation bifunctional protein PaaZ | 8.4 | 8.5 | 8.9 | 6.8 | 6.8 | 6.7 | 7.5 | 8.8 | 7.4 | 6.2 | 6.4 | 6.2 | 6.0 | 6.2 | 6.5 | **-2.46** | -0.64 | -0.04 |
| | | B5M07_01045 | Phenylacetic acid degradation protein | 8.2 | 7.6 | 7.2 | 5.2 | 4.7 | 4.8 | 5.4 | 7.3 | 6.2 | 4.9 | 5.0 | 5.2 | 4.7 | 5.3 | 5.3 | **-3.88** | **-1.58** | 0.14 |
| | | B5M07_01050 | Phenylacetate-CoA oxygenase subunit PaaJ | 7.8 | 7.5 | 7.7 | 3.7 | 4.3 | 4.1 | 5.1 | 6.9 | 5.0 | 3.6 | 4.3 | 4.1 | 3.5 | 4.0 | 4.4 | **-4.93** | **-2.13** | -0.10 |
| | | B5M07_01055 | Phenylacetate-CoA oxygenase subunit PaaI | 8.6 | 8.2 | 8.3 | 4.8 | 4.7 | 4.4 | 5.4 | 7.9 | 5.6 | 4.4 | 4.8 | 5.0 | 4.4 | 4.8 | 4.7 | **-5.07** | **-1.94** | -0.18 |
| | | B5M07_01060 | 1,2-Phenylacetyl-CoA epoxidase subunit B | 6.4 | 5.9 | 5.4 | 1.0 | 2.6 | 0.9 | 2.1 | 4.4 | 2.8 | 1.7 | 1.8 | 2.3 | 1.0 | 1.7 | 1.9 | **-6.15** | **-3.13** | -0.86 |
| | | B5M07_01065 | 1,2-Phenylacetyl-CoA epoxidase subunit A | 9.1 | 9.2 | 8.9 | 4.8 | 5.2 | 5.5 | 6.8 | 7.8 | 6.7 | 4.8 | 5.3 | 5.4 | 4.7 | 5.3 | 5.9 | **-5.29** | **-2.37** | 0.35 |
| | | B5M07_01820 | Phenylacetate--CoA ligase | 6.8 | 6.1 | 5.3 | 3.0 | 4.0 | 3.2 | 4.1 | 6.1 | 4.0 | 3.6 | 3.6 | 3.5 | 3.3 | 4.0 | 4.3 | **-3.95** | -1.43 | 0.52 |
| | | B5M07_06600 | 4-Hydroxyphenylpyruvate dioxygenase | 10.2 | 10.2 | 10.8 | 5.2 | 5.2 | 5.1 | 8.7 | 10.2 | 8.1 | 3.9 | 3.5 | 3.6 | 3.8 | 3.7 | 4.3 | **-7.00** | -1.31 | 0.72 |
| | | B5M07_17080 | Hypothetical protein | 3.6 | 3.7 | 4.5 | 3.3 | 1.8 | 1.2 | 2.7 | 3.9 | 2.7 | 1.6 | 2.4 | 0.8 | 1.2 | 1.2 | 2.2 | **-2.32** | -1.11 | -0.38 |
| 3 | Ribosome | B5M07_02120 | 30S ribosomal protein S15 | 3.2 | 4.3 | 3.0 | 6.0 | 5.5 | 6.0 | 3.2 | 4.0 | 4.8 | 4.8 | 3.4 | 4.2 | 4.9 | 3.7 | 2.9 | **3.23** | 0.88 | -0.19 |
| | | B5M07_05705 | 30S ribosomal protein S1 | 10.2 | 10.1 | 9.7 | 10.9 | 11.1 | 11.3 | 10.6 | 10.2 | 10.5 | 9.3 | 9.1 | 9.1 | 9.4 | 8.9 | 8.9 | **1.45** | 0.56 | -0.13 |
| | | B5M07_07690 | 30S ribosomal protein S6 | 4.1 | 6.1 | 5.1 | 5.8 | 6.1 | 6.4 | 4.2 | 3.9 | 4.0 | 5.6 | 5.6 | 4.7 | 5.5 | 4.0 | 3.7 | 1.03 | **-2.03** | -0.93 |
| | | B5M07_14705 | 50S ribosomal protein L17 | 2.7 | 4.0 | 2.6 | 5.6 | 5.6 | 5.8 | 4.6 | 5.1 | 4.5 | 5.1 | 4.5 | 4.7 | 4.3 | 3.7 | 3.1 | **3.89** | **2.70** | -1.36 |
| | | B5M07_14750 | 30S ribosomal protein S5 | 6.1 | 5.7 | 6.0 | 7.8 | 7.9 | 8.4 | 7.2 | 7.2 | 7.5 | 7.1 | 7.2 | 7.2 | 6.5 | 6.3 | 5.7 | **3.04** | **2.02** | **-1.30** |
| | | B5M07_14755 | 50S ribosomal protein L18 | 3.6 | 3.8 | 2.7 | 5.5 | 5.6 | 6.1 | 4.5 | 4.4 | 5.1 | 4.6 | 4.3 | 4.6 | 4.4 | 4.0 | 4.1 | **3.69** | **2.31** | -0.44 |

| | | | | | | | | | | | | | | | | | | | | | |
|---|---|---|---|---|---|---|---|---|---|---|---|---|---|---|---|---|---|---|---|---|---|
| | | B5M07_14760 | 50S ribosomal protein L6 | 5.4 | 5.3 | 5.4 | 6.5 | 7.0 | 7.5 | 5.7 | 6.1 | 6.4 | 6.2 | 6.3 | 6.2 | 5.7 | 5.4 | 5.3 | **2.42** | 1.15 | **-1.10** |
| | | B5M07_14765 | 30S ribosomal protein S8 | 4.2 | 3.4 | 4.4 | 5.2 | 5.0 | 5.5 | 3.1 | 3.3 | 3.7 | 4.7 | 4.2 | 4.6 | 3.9 | 3.0 | 3.3 | **1.71** | -1.04 | **-1.52** |
| | | B5M07_14785 | 50S ribosomal protein L14 | 3.2 | 2.7 | 3.6 | 4.3 | 4.6 | 4.8 | 2.9 | 2.9 | 3.4 | 3.9 | 4.0 | 3.4 | 3.6 | 1.9 | 3.3 | **2.05** | -0.22 | -0.95 |
| | | B5M07_14790 | 30S ribosomal protein S17 | 1.8 | 4.1 | 3.1 | 4.1 | 4.3 | 4.3 | 1.3 | 1.4 | 1.2 | 2.8 | 2.7 | 2.8 | 3.6 | 2.6 | 2.5 | 1.17 | **-5.59** | 0.45 |
| | | B5M07_14865 | 50S ribosomal protein L4 | 5.8 | 5.4 | 5.0 | 6.8 | 7.2 | 7.5 | 6.0 | 5.7 | 5.8 | 6.4 | 6.2 | 6.0 | 6.2 | 5.6 | 5.5 | **2.51** | 0.66 | -0.56 |
| | | B5M07_14890 | 30S ribosomal protein S7 | 3.4 | 5.5 | 4.5 | 7.0 | 7.1 | 7.8 | 5.5 | 3.9 | 4.7 | 5.8 | 5.4 | 5.3 | 5.4 | 5.4 | 4.9 | **3.50** | 0.15 | -0.37 |
| | | B5M07_14895 | 30S ribosomal protein S12 | 4.5 | 3.9 | 3.2 | 5.8 | 5.5 | 6.5 | 5.0 | 3.3 | 4.2 | 5.3 | 4.5 | 3.7 | 5.2 | 4.3 | 3.9 | **2.96** | 0.68 | -0.04 |
| | | B5M07_14940 | 50S ribosomal protein L11 | 6.4 | 6.7 | 6.5 | 7.9 | 7.7 | 7.7 | 6.4 | 6.0 | 5.9 | 7.1 | 6.9 | 6.6 | 7.0 | 6.6 | 5.9 | **1.70** | -0.65 | -0.38 |
| | | B5M07_15210 | 50S ribosomal protein L19 | 5.7 | 5.8 | 4.8 | 6.7 | 6.7 | 6.7 | 6.1 | 5.9 | 5.8 | 5.7 | 5.2 | 4.8 | 5.8 | 5.5 | 5.1 | **1.61** | 0.56 | 0.33 |
| 4 | Oxidative phospho-rylation | B5M07_13780 | ATP synthase F1 subunit epsilon | 5.2 | 4.4 | 3.6 | 5.9 | 5.7 | 6.0 | 5.6 | 5.9 | 5.9 | 5.7 | 5.5 | 5.7 | 5.6 | 5.3 | 4.7 | **2.04** | **1.94** | -0.55 |
| | | B5M07_13785 | F0F1 ATP synthase subunit beta | 7.6 | 6.9 | 7.4 | 8.5 | 8.7 | 8.9 | 8.2 | 8.5 | 8.6 | 8.8 | 8.6 | 8.4 | 8.3 | 8.5 | 8.2 | **1.99** | **1.64** | -0.37 |
| | | B5M07_13790 | F0F1 ATP synthase subunit gamma | 6.2 | 5.7 | 5.3 | 6.7 | 6.9 | 6.9 | 6.6 | 6.9 | 6.7 | 6.9 | 6.9 | 6.6 | 6.8 | 6.5 | 6.4 | **1.52** | **1.42** | -0.26 |
| | | B5M07_13795 | F0F1 ATP synthase subunit alpha | 7.6 | 6.7 | 7.4 | 8.4 | 8.3 | 8.8 | 7.6 | 7.6 | 7.9 | 8.8 | 8.6 | 8.4 | 8.7 | 8.5 | 8.6 | **1.75** | 0.64 | -0.02 |
| | | B5M07_15125 | ATP F0F1 synthase subunit B | 4.2 | 3.6 | 3.2 | 5.3 | 4.4 | 5.0 | 4.1 | 4.6 | 4.8 | 5.8 | 4.5 | 4.4 | 5.3 | 5.0 | 4.4 | **2.04** | 1.45 | -0.19 |
| | | B5M07_15135 | F0F1 ATP synthase subunit C | 2.7 | 2.1 | 2.0 | 3.3 | 3.9 | 3.9 | 2.4 | 4.3 | 2.7 | 4.7 | 3.7 | 4.1 | 4.4 | 3.7 | 2.6 | **3.60** | 3.13 | -0.64 |

[a] Clusters of differentially expressed genes presented in Fig. 1d
[b] Based on NCBI Prokaryotic Genome Annotation Pipeline
[c] Values in bold are statistically significant (|fold change|>2 and adjusted *P*-value ≤ 0.05)

**Table S5.** ***Sulfitobacter*** **D7 expression of flagellar genes presented in Fig. 2.**

| Category | Gene | Locus tag | Normalized expression values (log transformed) (triplicate samples for each treatment) | | | | | | | | | | | | | | | Differential expression[a] Fold change ($log_2$) | | |
|---|---|---|---|---|---|---|---|---|---|---|---|---|---|---|---|---|---|---|---|---|
| | | | Exp-CM | | | Exp-CM +DMSP | | | Stat-CM | | | MM | | | MM +DMSP | | | Exp-CM +DMSP vs. Exp-CM | Stat-CM vs. Exp-CM | MM +DMSP vs. MM |
| C ring, switch complex | *FliG* | B5M07_04650 | 3.7 | 5.3 | 5.4 | 6.1 | 5.5 | 6.0 | 6.3 | 5.1 | 6.1 | 6.9 | 7.1 | 7.1 | 6.7 | 6.7 | 6.6 | **1.37** | **1.46** | -0.51 |
| | *FliM* | B5M07_04680 | 7.2 | 7.3 | 7.6 | 7.4 | 7.4 | 6.9 | 8.2 | 7.2 | 7.9 | 8.5 | 8.7 | 8.8 | 8.1 | 8.4 | 8.6 | -0.2 | 0.7 | -0.45 |
| | *FliN* | B5M07_04660 | 4.5 | 4.0 | 3.2 | 5.7 | 5.3 | 5.4 | 5.5 | 4.0 | 5.2 | 6.4 | 6.8 | 6.3 | 5.8 | 6.1 | 6.0 | **2.74** | **2.11** | -0.72 |
| Rod | *FlgB* | B5M07_04740 | 5.5 | 6.2 | 5.8 | 5.5 | 5.3 | 5.0 | 6.2 | 5.5 | 6.3 | 7.2 | 6.8 | 6.5 | 7.0 | 6.6 | 6.8 | -0.91 | 0.28 | -0.1 |
| | *FlgC* | B5M07_04735 | 3.4 | 3.9 | 5.2 | 4.4 | 4.5 | 4.6 | 5.0 | 3.6 | 5.1 | 4.9 | 5.3 | 5.1 | 5.0 | 4.8 | 5.0 | 0.17 | 0.52 | -0.2 |
| | *FlgF* | B5M07_04590 | 5.1 | 5.3 | 5.1 | 6.4 | 6.6 | 5.6 | 7.5 | 5.7 | 7.2 | 7.3 | 7.2 | 7.0 | 6.7 | 7.0 | 7.3 | **1.76** | **2.84** | -0.24 |
| | *FlgG* | B5M07_04585 | 5.3 | 4.6 | 4.7 | 6.4 | 5.9 | 5.7 | 6.6 | 5.1 | 6.7 | 7.3 | 7.4 | 7.3 | 6.4 | 6.8 | 7.1 | **1.93** | **2.38** | -0.7 |
| Rod-associated | *FliE* | B5M07_04730 | 3.4 | 1.9 | 1.9 | 2.5 | 1.8 | 2.3 | 3.8 | 2.5 | 3.5 | 3.9 | 4.4 | 4.7 | 3.8 | 3.6 | 3.7 | -0.28 | 1.64 | -0.92 |
| | *FliL* | B5M07_04675 | 7.5 | 6.8 | 6.7 | 6.4 | 6.3 | 6.2 | 7.7 | 7.6 | 8.1 | 8.2 | 7.8 | 8.2 | 8.2 | 8.0 | 8.3 | **-1.12** | **1.09** | 0.08 |
| L ring | *FlgH* | B5M07_04575 | 2.8 | 2.8 | 2.7 | 4.8 | 4.8 | 4.6 | 5.0 | 3.5 | 4.1 | 5.9 | 6.1 | 5.8 | 5.0 | 5.3 | 5.2 | **5.43** | **4.89** | **-1.1** |
| P ring | *FlgI* | B5M07_04615 | 4.1 | 5.6 | 5.2 | 7.0 | 7.0 | 7.0 | 7.0 | 5.9 | 7.1 | 7.3 | 7.7 | 7.7 | 7.1 | 7.0 | 7.0 | **3.02** | **2.73** | -0.75 |
| P ring chaperon | *FlgA* | B5M07_04580 | 4.2 | 3.2 | 4.6 | 5.5 | 5.3 | 4.6 | 5.5 | 4.4 | 5.3 | 6.1 | 6.4 | 6.3 | 5.3 | 5.7 | 5.5 | **1.81** | **1.77** | -1.02 |
| MS ring | *FliF* | B5M07_04670 | 5.7 | 5.5 | 6.6 | 6.9 | 6.9 | 6.5 | 7.0 | 6.4 | 7.2 | 8.5 | 8.5 | 8.3 | 8.1 | 8.4 | 8.3 | **1.22** | **1.41** | -0.18 |
| Ion channels (PMF) | *MotA* | B5M07_04535 | 5.0 | 6.0 | 4.7 | 6.2 | 6.0 | 5.9 | 6.4 | 5.4 | 6.9 | 7.5 | 7.8 | 7.7 | 7.2 | 7.4 | 7.4 | 1.13 | **1.68** | -0.44 |
| | *MotB* | B5M07_04550 | 6.3 | 5.0 | 5.3 | 6.3 | 6.1 | 6.2 | 6.9 | 6.4 | 6.5 | 7.2 | 7.4 | 7.2 | 6.4 | 6.6 | 6.6 | 0.79 | **1.38** | -0.96 |
| | *MotB* | B5M07_13825 | 5.5 | 4.8 | 5.8 | 6.6 | 5.9 | 6.5 | 6.3 | 5.4 | 6.4 | 7.0 | 7.0 | 7.0 | 6.9 | 6.9 | 6.6 | **1.43** | 1.07 | -0.24 |
| T3SS | *FlhA* | B5M07_04705 | 0.0 | 0.0 | -0.8 | -0.5 | 0.6 | -1.2 | -0.4 | -1.0 | -1.1 | -1.0 | -1.3 | -1.4 | -0.7 | -1.3 | -1.2 | -0.4 | NA | NA |
| | *FlhB* | B5M07_04695 | 3.6 | 2.2 | 2.1 | 1.8 | 2.5 | 1.3 | 2.2 | 2.4 | 2.3 | 2.3 | 3.0 | 2.7 | 2.6 | 2.9 | 2.7 | -1.62 | -0.81 | 0.07 |
| | *FliQ* | B5M07_04725 | 1.9 | 1.8 | 1.0 | 1.4 | 0.6 | 1.5 | 0.7 | 1.8 | 1.2 | 2.8 | 3.3 | 2.5 | 2.7 | 2.2 | 1.5 | -0.56 | NA | NA |
| | *FliP* | B5M07_04655 | 4.2 | 5.1 | 4.9 | 6.8 | 6.5 | 6.3 | 6.1 | 4.8 | 6.3 | 7.4 | 7.9 | 7.5 | 7.0 | 7.1 | 7.1 | **2.84** | **1.94** | -0.73 |
| | *FliR* | B5M07_04700 | 0.0 | 0.0 | 0.0 | 0.0 | 0.0 | 0.0 | 0.0 | 0.0 | 0.0 | 0.0 | 0.0 | 0.0 | 0.0 | 0.0 | 0.0 | NA | NA | NA |
| ATPase complex | *FliH* | B5M07_04665 | 4.1 | 3.3 | 4.0 | 5.6 | 5.0 | 5.2 | 5.3 | 4.7 | 5.1 | 6.8 | 6.9 | 6.9 | 6.4 | 6.2 | 6.2 | **2.99** | **2.7** | -0.82 |
| | *FliI* | B5M07_04745 | 6.6 | 6.0 | 6.3 | 6.0 | 5.4 | 5.8 | 6.6 | 6.5 | 6.5 | 7.0 | 6.6 | 6.9 | 6.4 | 6.4 | 6.3 | -0.8 | 0.31 | -0.67 |
| Filament | *FliC* | B5M07_04630 | 11.6 | 11.6 | 11.3 | 10.7 | 10.6 | 10.5 | 11.5 | 11.4 | 11.5 | 10.2 | 10.3 | 10.2 | 10.2 | 10.3 | 10.5 | **-1.28** | -0.08 | 0.18 |

| | | | | | | | | | | | | | | | | | | | |
|---|---|---|---|---|---|---|---|---|---|---|---|---|---|---|---|---|---|---|---|---|
| Hook control | *FliK* | B5M07_04560 | 6.0 | 6.2 | 6.5 | 7.4 | 7.5 | 7.3 | 8.2 | 6.6 | 7.7 | 8.1 | 8.4 | 8.4 | 7.6 | 7.6 | 7.6 | **1.78** | **2.19** | -0.92 |
| | *FlgD* | B5M07_04565 | 3.6 | 3.5 | 4.7 | 5.4 | 4.8 | 5.1 | 5.5 | 4.6 | 5.2 | 5.9 | 6.4 | 6.1 | 5.1 | 5.4 | 5.6 | **1.71** | **1.76** | -1.04 |
| Hook | *FlgE* | B5M07_04600 | 7.1 | 7.0 | 7.0 | 8.3 | 8.3 | 8.0 | 8.8 | 7.6 | 8.7 | 8.9 | 9.1 | 8.8 | 8.9 | 9.4 | 9.4 | **1.71** | **2.17** | 0.39 |
| Hook-filament junction | *FlgL* | B5M07_04610 | 5.1 | 3.8 | 4.6 | 6.5 | 6.5 | 6.7 | 6.8 | 6.0 | 6.4 | 7.2 | 7.6 | 7.2 | 6.8 | 6.6 | 6.9 | **3.44** | **3.28** | -0.73 |
| | *FlgK* | B5M07_04605 | 5.8 | 6.1 | 6.3 | 8.3 | 8.1 | 8.1 | 8.5 | 7.5 | 8.5 | 9.1 | 9.2 | 9.2 | 8.6 | 8.9 | 9.1 | **3.39** | **3.5** | -0.4 |

[a] Values in bold are statistically significant (|fold change|>2 and adjusted *P*-value ≤ 0.05). NA – could not calculate due to low detection limit

**<u>Table S6.</u>** ***Sulfitobacter*** **D7 expression of DE transport genes.**

File attached.

**Table S7.** ***Sulfitobacter*** **D7 expression of DMSP transport and metabolism genes.**

| Category | Gene | Locus tag | Normalized expression values (log transformed) (triplicate samples for each treatment) | | | | | | | | | | | | | | | Differential expression[a] Fold change ($log_2$) | | |
|---|---|---|---|---|---|---|---|---|---|---|---|---|---|---|---|---|---|---|---|---|
| | | | Exp-CM | | | Exp-CM +DMSP | | | Stat-CM | | | MM | | | MM +DMSP | | | Exp-CM +DMSP vs. Exp-CM | Stat-CM vs. Exp-CM | MM +DMSP vs. MM |
| DMSP transporters | *BCCT* | B5M07_05880 | 6.2 | 5.8 | 5.6 | 6.8 | 7.7 | 8.0 | 6.9 | 6.5 | 7.4 | 5.6 | 6.7 | 6.9 | 6.0 | 6.5 | 6.9 | **2.48** | **1.69** | -0.09 |
| | *BCCT* | B5M07_09045 | 7.1 | 7.2 | 6.7 | 8.0 | 8.1 | 8.1 | 7.9 | 7.5 | 7.8 | 7.3 | 7.6 | 7.7 | 7.3 | 7.7 | 7.6 | **1.5** | **1.08** | 0.02 |
| DMSP demethylation enzymes | *DmdA* | B5M07_09030 | 5.2 | 4.7 | 5.2 | 8.0 | 7.8 | 7.9 | 7.0 | 6.6 | 7.6 | 4.2 | 3.9 | 3.9 | 4.7 | 4.2 | 4.2 | **3.86** | **2.87** | 0.65 |
| | *DmdB* | B5M07_09700 | 4.8 | 4.2 | 5.6 | 5.5 | 5.1 | 5.1 | 5.3 | 4.6 | 5.3 | 5.8 | 6.1 | 5.6 | 5.6 | 5.5 | 5.8 | 0.3 | 0.09 | -0.28 |
| | *$DmdC_1$* | B5M07_13415 | 10.0 | 9.6 | 10.9 | 9.7 | 9.4 | 10.7 | 8.4 | 8.6 | 8.2 | 4.7 | 5.4 | 5.4 | 4.8 | 4.8 | 5.1 | -0.36 | **-2.46** | -0.82 |
| | *$DmdC_2$* | B5M07_13985 | 7.2 | 6.9 | 7.5 | 7.2 | 6.9 | 7.3 | 8.0 | 7.4 | 8.0 | 5.4 | 5.1 | 5.0 | 5.3 | 4.8 | 5.6 | -0.05 | 0.78 | 0.17 |
| | *DmdD* | B5M07_06160 | 4.8 | 5.0 | 5.4 | 4.7 | 4.9 | 5.6 | 5.8 | 5.8 | 6.0 | 3.2 | 3.2 | 3.1 | 3.5 | 3.0 | 3.5 | 0.06 | 0.98 | 0.27 |

[a] Values in bold are statistically significant (|fold change|>2 and adjusted *P*-value ≤ 0.05)

### Table S8. *Sulfitobacter* D7 expression of benzoate transport and metabolism genes presented in Fig. 3a.

| Category | Gene | Locus tag | Normalized expression values (log transformed) (triplicate samples for each treatment) | | | | | | | | | | | | | | | Differential expression[a] Fold change ($log_2$) | | |
|---|---|---|---|---|---|---|---|---|---|---|---|---|---|---|---|---|---|---|---|---|
| | | | Exp-CM | | | Exp-CM +DMSP | | | Stat-CM | | | MM | | | MM +DMSP | | | Exp-CM +DMSP vs. Exp-CM | Stat-CM vs. Exp-CM | MM +DMSP vs. MM |
| Transporter | *BenE* | B5M07_03645 | 4.1 | 3.2 | 3.4 | 3.0 | 3.9 | 3.8 | 3.4 | 4.2 | 3.5 | 2.2 | 2.5 | 2.0 | 1.2 | 2.4 | 1.5 | -0.05 | 0.1 | -1 |
| Transcription | *BenM* | B5M07_19060 | 5.3 | 5.5 | 6.4 | 7.5 | 7.2 | 7.4 | 6.6 | 4.9 | 6.3 | 4.7 | 4.9 | 4.6 | 4.7 | 4.9 | 4.7 | **2.07** | 0.43 | 0.05 |
| Benzoate catabolism enzymes | *BenA* | B5M07_19105 | 9.3 | 10.2 | 10.9 | 12.1 | 12.1 | 12.4 | 12.2 | 9.6 | 11.7 | 6.9 | 6.8 | 6.7 | 7.0 | 7.8 | 8.3 | **2.36** | 1.56 | **2.19** |
| | *BenB* | B5M07_19100 | 5.5 | 7.5 | 8.0 | 9.8 | 9.8 | 10.0 | 9.7 | 7.3 | 9.3 | 4.5 | 4.1 | 4.0 | 4.7 | 5.5 | 6.0 | **3.19** | **2.24** | **2.7** |
| | *BenC* | B5M07_19095 | 4.9 | 6.5 | 8.1 | 10.2 | 10.2 | 10.5 | 10.3 | 6.5 | 9.1 | 4.6 | 4.6 | 4.3 | 4.5 | 5.3 | 6.0 | **4.01** | 2.88 | 1.84 |
| | *BenD* | B5M07_19090 | 4.5 | 4.6 | 6.8 | 9.1 | 9.3 | 9.7 | 9.2 | 4.5 | 7.4 | 3.3 | 3.5 | 3.6 | 3.8 | 4.7 | 4.2 | **4.53** | 3 | 1.73 |
| | *CatA* | B5M07_19055 | 9.7 | 11.5 | 11.0 | 11.5 | 11.7 | 11.7 | 11.0 | 8.4 | 10.6 | 6.5 | 6.3 | 6.3 | 6.5 | 6.7 | 6.7 | 0.84 | -0.7 | 0.58 |
| | *CatB* | B5M07_19110 | 9.2 | 11.1 | 10.5 | 9.6 | 9.8 | 9.4 | 9.7 | 8.3 | 9.8 | 6.0 | 5.8 | 5.8 | 6.0 | 6.2 | 6.6 | -1.16 | -1.37 | 0.8 |
| | *PcaD* | B5M07_19210 | 1.7 | 2.2 | 1.6 | 2.3 | 1.0 | 0.4 | 2.4 | 0.7 | 1.7 | 1.6 | 2.0 | 1.2 | 1.1 | 1.6 | 2.2 | -0.76 | NA | NA |
| | *PcaI* | B5M07_19140 | 2.4 | 4.5 | 4.7 | 2.7 | 3.0 | 3.0 | 3.1 | 2.3 | 3.4 | 1.9 | 2.0 | 2.7 | 2.3 | 1.9 | 2.0 | **-1.86** | -1.7 | -0.31 |
| | *PcaJ* | B5M07_19145 | 3.3 | 2.6 | 3.2 | 1.0 | 2.1 | 2.7 | 3.1 | 2.7 | 2.9 | 2.5 | 2.0 | 2.0 | 2.1 | 1.9 | 2.2 | -1.63 | -0.32 | -0.23 |
| | *PcaF* | B5M07_19150 | 4.8 | 5.3 | 6.5 | 4.7 | 4.3 | 4.9 | 5.4 | 4.9 | 5.5 | 4.2 | 4.2 | 4.1 | 3.9 | 3.6 | 3.7 | **-1.54** | -0.63 | -0.63 |

[a] Values in bold are statistically significant (|fold change|>2 and adjusted *P*-value ≤ 0.05). NA – could not calculate due to low detection limit

### <u>Table S9.</u> Presence of benzoate degradation and transport genes in genomes of phytoplankton-associated bacteria.

| Bacterial Strain | Benzoate-related genes | | | | | | | | | | | | | Genome accession number[a] | References |
|---|---|---|---|---|---|---|---|---|---|---|---|---|---|---|---|
| | Degradation pathway[b] | | | | | | | | | | | | | | |
| | Transport | | | | | Catechol | | | Proto-catecuate | | Benzoyl-CoA | | | | |
| | *BenK1* | *BenK2* | *BenE1* | *BenE2* | *BenF* | *BenA* | *BenB* | *BenC* | *BphA* | *PobA* | CoA-ligase | *BoxA* | *BoxB* | | |
| ***Sulfitobacter* D7** | | | + | + | | + | + | + | Partial | Partial | + | | | 1968541 | (Barak-Gavish *et al.*, 2018) (3) |
| ***Sulfitobacter* sp. CB2047** | | | + | + | | + | + | + | Partial | + | + | Partial | | 1525218 | (Ankrah *et al.*, 2014) (4) |
| *Sulfitobacter pseudonitzschiae* F5 | Partial | Partial | | | | + | | + | Partial | + | + | + | + | GCF_014805345.1 | (Amin *et al.*, 2015; Fei *et al.*, 2020) (5, 6) |
| *Sulfitobacter geojensis* EhN01 | | Partial | | | | + | | + | Partial | + | + | + | + | GCF_001650795.1 | (Orata *et al.*, 2016) (7) |
| *Sulfitobacter pontiacus* EhN02 | | + | | | | Partial | | + | Partial | + | + | | | GCF_001650835.1 | (Orata *et al.*, 2016) (7) |
| *Sulfitobacter* sp. EhC04 | | | | | | + | + | + | + | + | + | + | + | 1849168 | (Rosana *et al.*, 2016) (8) |
| ***Rhodobacteraceae* bacterium EhC02** | | | + | + | | + | | + | + | + | + | Partial | | 1849170 | (Rosana *et al.*, 2016) (8) |
| ***Roseovarius indicus* EhC03** | | + | + | + | | + | | + | + | + | + | + | + | GCF_001650975.1 | (Rosana *et al.*, 2016) (8) |
| ***Ruegeria pomeroyi* DSS-3** | | | + | + | | Partial | Partial | Partial | Partial | + | + | + | + | 246200 | (Durham *et al.*, 2015; Landa *et al.*, 2017) (9, 10) |
| *Ruegeria* sp. R11 | | | + | + | | Partial | | Partial | Partial | + | + | Partial | | 439497 | (Mayers *et al.*, 2016) (11) |
| *Ruegeria* sp. TM1040 | | | + | + | | + | | + | Partial | + | + | Partial | | 292414 | (Miller *et al.*, 2004; Miller and Belas, 2006) (12, 13) |
| *Phaeobacter inhibens* DSM 17395 | | + | + | + | | + | | + | Partial | + | + | Partial | | 391619 | (Seyedsayamdost *et al.*, 2011; Segev *et al.*, 2016) (14, 15) |
| *Dinoroseobacter shibae* DFL-12 | | | | | | Partial | | Partial | Partial | + | + | Partial | | 398580 | (Wang *et al.*, 2014, 2015) (16, 17) |
| *Jannaschia* sp. EhC01 | | | | | | + | | + | + | + | + | Partial | + | 1849359 | (Rosana *et al.*, 2016) (8) |
| *Erythrobacter* sp. EhN03 | Partial | Partial | | | | Partial | | | | | + | | | 1849358 | (Orata *et al.*, 2016) (7) |
| ***Sphingomonadales* bacterium EhC05** | + | + | | | | + | | + | + | + | + | | | 1849171 | (Rosana *et al.*, 2016) (8) |
| ***Marinobacter adhaerens*** | | + | + | + | | + | + | + | | | + | + | | 1033846 | (Amin *et al.*, 2009; Sonnenschein *et al.*, 2012) (18, 19) |

| | | | | | | | | | | | | | | | |
|---|---|---|---|---|---|---|---|---|---|---|---|---|---|---|---|
| ***Marinobacter algicola*** | + | + | + | + | | + | + | + | | | + | + | | 236100 | (Amin *et al.*, 2009; Green *et al.*, 2015) (18, 20) |
| ***Marinobacter* sp. EhC06** | | + | | | | + | + | + | | | + | + | | 1849169 | (Rosana *et al.*, 2016) (8) |
| *Marinobacter* sp. EhN04 | | | | + | | + | | + | | | + | + | | 1849167 | (Orata *et al.*, 2016) (7) |
| *Pseudoalteromonas piscicida* A757 | + | Partial | + | + | | Partial | | + | | | + | + | | GCF_004103285.1 | (Harvey *et al.*, 2016) (21) |
| *Balneola* sp. EhC07 | | Partial | | | | | | + | | + | + | + | | 1849360 | (Rosana *et al.*, 2016) (8) |

[a]Taxonomy ID or RefSeq assembly accession

[b]Bacterial benzoate degradation pathways are elaborated in Fig. S3

Colored tiles depict the presence of a transporter or the presence of all the examined query genes in each degradation pathway

Bacterial strains highlighted in **bold** harbor genes for both transport and catabolism and thus can utilize benzoate

The query genes are listed in Table S10

**<u>Table S10.</u> Benzoate degradation and transport genes used as queries for BLASTp against genomes of phytoplankton-associated bacteria.**

| | | Gene name | Function | Reaction | Accession number | Organism | Reference |
|---|---|---|---|---|---|---|---|
| Transport | | *$BenK_1$* | Transport | | AAC46425.1 | *Acinetobacter* sp. strain ADP1 | Collier *et al.*, 1997 (22) |
| | | *$BenK_2$* | Transport | | AAN68773.1 | *Pseudomonas putida* KT2440 | Nishikawa *et al.*, 2008 (23) |
| | | *$BenE_1$* | Transport | | AAN67649.1 | *Pseudomonas putida* KT2440 | |
| | | *$BenE_2$* | Transport | | AAN68775.1 | *Pseudomonas putida* KT2440 | |
| | | *BenF* | Transport | | AAN68776.1 | *Pseudomonas putida* KT2440 | |
| Degradation pathway | Catechol | *BenA* | Benzoate 1,2-dioxygenase subunit alpha | Benzoate → *cis*-1,2-dihydroxybenzoate | AAC46436.2 | *Acinetobacter* sp. strain ADP1 | Collier *et al.*, 1998 (24) |
| | | *BenB* | Benzoate 1,2-dioxygenase subunit beta | | AAC46437.1 | *Acinetobacter* sp. strain ADP1 | |
| | | *BenC* | Benzoate 1,2-dioxygenase electron transfer component | | AAC46438.1 | *Acinetobacter* sp. strain ADP1 | |
| | Proto-catecuate | *BphA* | Benzoate 4-monooxygenase | Benzoate → 4-OH-Benzoate | P17549.1 | *Aspergillus niger* | van Gorcom *et al.*, 1990 (25) |
| | | *PobA* | 4-hydroxybenzoate 3-monooxygenase | 4-OH-Benzoate→ protocatecuate | WP_004926674.1 | *Acinetobacter* sp. strain ADP1 | Brzostowicz *et al.*, 2003 (26) |
| | Benzoyl-CoA | CoA-ligase | Benzoate CoA-ligase | Benzoate→ Benzoyl-CoA | AAN39371.1 | *Azoarcus evansii* | Mohamed *et al.*, 2001 (27)<br>Gescher *et al.*, 2002 (28) |
| | | *BoxA* | Benzoyl-CoA oxygenase component A | Benzoyl-CoA→ 2,3-epoxy-2,3-dihydrobenzoyl-CoA | AAN39377.1 | *Azoarcus evansii* | Gescher *et al.*, 2002 (28) |
| | | *BoxB* | Benzoyl-CoA oxygenase component B | | AAN39376.1 | *Azoarcus evansii* | |

All genes were experimentally validated, except *BenC*

**Table S11.** **Statistical significance of differences in algal and bacterial growth in the comparisons between the treatments presented in Fig. 5.**

| Parameter | Treatments comparison[a] | *P*-value[b] |
|---|---|---|
| Algal growth | Ehux vs. Ehux+DMSP | 0.47 |
| | Ehux vs. Ehux+Benzoate | 1.00 |
| | Ehux vs. Ehux+Benzoate+DMSP | 0.21 |
| | Ehux vs. Ehux+D7 | **0.01** |
| | Ehux vs. Ehux+D7+DMSP | **<.0001** |
| | Ehux vs. Ehux+D7+Benzoate | 0.05 |
| | Ehux vs. Ehux+D7+Benzoate+DMSP | 1.00 |
| | Ehux+DMSP vs. Ehux+Benzoate | 0.28 |
| | Ehux+DMSP vs. Ehux+Benzoate+DMSP | 1.00 |
| | Ehux+DMSP vs. Ehux+D7 | **<.0001** |
| | Ehux+DMSP vs. Ehux+D7+DMSP | **<.0001** |
| | Ehux+DMSP vs. Ehux+D7+Benzoate | **<.0001** |
| | Ehux+DMSP vs. Ehux+D7+Benzoate+DMSP | 0.21 |
| | Ehux+Benzoate vs. Ehux+Benzoate+DMSP | 0.11 |
| | Ehux+Benzoate vs. Ehux+D7 | **0.03** |
| | Ehux+Benzoate vs. Ehux+D7+DMSP | **<.0001** |
| | Ehux+Benzoate vs. Ehux+D7+Benzoate | 0.12 |
| | Ehux+Benzoate vs. Ehux+D7+Benzoate+DMSP | 1.00 |
| | Ehux+Benzoate+DMSP vs. Ehux+D7 | **<.0001** |
| | Ehux+Benzoate+DMSP vs. Ehux+D7+DMSP | **<.0001** |
| | Ehux+Benzoate+DMSP vs. Ehux+D7+Benzoate | **<.0001** |
| | Ehux+Benzoate+DMSP vs. Ehux+D7+Benzoate+DMSP | 0.07 |
| | Ehux+D7 vs. Ehux+D7+DMSP | **<.0001** |
| | Ehux+D7 vs. Ehux+D7+Benzaote | 1.00 |
| | Ehux+D7 vs. Ehux+D7+Benzoate+DMSP | **0.04** |
| | Ehux+D7+DMSP vs. Ehux+D7+Benzoate | **<.0001** |
| | Ehux+D7+DMSP vs. Ehux+D7+Benzoate+DMSP | **<.0001** |
| | Ehux+D7+Benzoate vs. Ehux+D7+Benzoate+DMSP | 0.17 |
| Bacterial growth | Ehux+D7 vs. Ehux+D7+DMSP | 0.46 |
| | Ehux+D7 vs. Ehux+D7+Benzoate | **0.01** |
| | Ehux+D7 vs. Ehux+D7+Benzoate+DMSP | **0.04** |
| | Ehux+D7+DMSP vs. Ehux+D7+Benzoate | 0.28 |

| | |
|---|---|
| Ehux+D7+DMSP vs. Ehux+D7+Benzoate+DMSP | 0.58 |
| Ehux+D7+Benzoate vs. Ehux+D7+Benzoate+DMSP | 0.96 |

[a] Ehux, *E. huxleyi*; D7, *Sulfitobacter* D7

[b] Values in bold are statistically significant (*P*-value < 0.05)

**Table S12.** Number of sequencing reads of *Sulfitobacter* D7 transcriptome experiment.

| Treatment | Replicate | Raw reads | Reads after adapter trimming and quality control | Reads aligned concordantly to *Sulfitobacter* D7 genome |
|---|---|---|---|---|
| Exp-CM | 1 | 767,618 | 745,975 | 176,676 |
| | 2 | 922,968 | 906,039 | 191,517 |
| | 3 | 1,151,749 | 1,129,112 | 226,304 |
| Exp-CM+DMSP | 1 | 1,534,356 | 1,480,621 | 685,162 |
| | 2 | 1,589,860 | 1,571,506 | 759,918 |
| | 3 | 2,407,636 | 2,369,101 | 1,185,338 |
| Stat-CM | 1 | 1,322,423 | 1,297,304 | 464,970 |
| | 2 | 839,261 | 821,289 | 314,033 |
| | 3 | 1,313,839 | 1,297,430 | 512,039 |
| MM | 1 | 6,120,691 | 5,850,724 | 1,222,165 |
| | 2 | 12,093,273 | 11,808,396 | 1,151,032 |
| | 3 | 10,670,288 | 10,410,612 | 1,417,129 |
| MM+DMSP | 1 | 9,290,554 | 9,017,929 | 1,934,490 |
| | 2 | 7,948,137 | 7,804,118 | 1,114,609 |
| | 3 | 6,528,722 | 6,396,058 | 895,160 |

**SI References**


1. T. Hulsen, J. de Vlieg, W. Alkema, BioVenn – a web application for the comparison and visualization of biological lists using area-proportional Venn diagrams. *BMC Genomics* **9**, 488 (2008).
2. G. Fuchs, M. Boll, J. Heider, Microbial degradation of aromatic compounds — from one strategy to four. *Nat. Rev. Microbiol.* **9**, 803–816 (2011).
3. N. Barak-Gavish, *et al.*, Bacterial virulence against an oceanic bloom-forming phytoplankter is mediated by algal DMSP. *Sci. Adv.* **4**, eaau5716 (2018).
4. N. Y. D. Ankrah, T. Lane, C. R. Budinoff, M. K. Hadden, A. Buchan, Draft Genome sequence of *Sulfitobacter* sp. CB2047, a member of the *Roseobacter* clade of marine bacteria, isolated from an *Emiliania huxleyi* Bloom. *Genome Announc.* **2**, e01125-14 (2014).
5. S. A. Amin, *et al.*, Interaction and signalling between a cosmopolitan phytoplankton and associated bacteria. *Nature* **522**, 98–101 (2015).
6. C. Fei, *et al.*, Quorum sensing regulates 'swim-or-stick' lifestyle in the phycosphere. *Environ. Microbiol.* **22**, 4761–4778 (2020).
7. F. D. Orata, *et al.*, Draft Genome Sequences of Four Bacterial Strains Isolated from a Polymicrobial Culture of Naked (N-Type) *Emiliania huxleyi* CCMP1516. *Genome Announc.* **4**, 9–10 (2016).
8. A. R. R. Rosana, *et al.*, Draft Genome Sequences of Seven Bacterial Strains Isolated from a Polymicrobial Culture of Coccolith-Bearing (C-Type) *Emiliania huxleyi* M217. *Genome Announc.* **4**, 9–10 (2016).
9. B. P. Durham, *et al.*, Cryptic carbon and sulfur cycling between surface ocean plankton. *Proc. Natl. Acad. Sci.* **112**, 453–457 (2015).
10. M. Landa, A. S. Burns, S. J. Roth, M. A. Moran, Bacterial transcriptome remodeling during sequential co-culture with a marine dinoflagellate and diatom. *ISME J.* **11**, 2677–2690 (2017).
11. T. J. Mayers, A. R. Bramucci, K. M. Yakimovich, R. J. Case, A Bacterial Pathogen Displaying Temperature-Enhanced Virulence of the Microalga *Emiliania huxleyi*. *Front. Microbiol.* **7**, 892 (2016).
12. T. R. Miller, K. Hnilicka, A. Dziedzic, P. Desplats, R. Belas, Chemotaxis of *Silicibacter* sp. strain TM1040 toward dinoflagellate products. *Appl. Environ. Microbiol.* **70**, 4692–4701 (2004).
13. T. R. Miller, R. Belas, Motility is involved in *Silicibacter* sp. TM1040 interaction with dinoflagellates. *Environ. Microbiol.* **8**, 1648–1659 (2006).
14. M. R. Seyedsayamdost, R. J. Case, R. Kolter, J. Clardy, The Jekyll-and-Hyde chemistry of *Phaeobacter gallaeciensis*. *Nat. Chem.* **3**, 331–5 (2011).
15. E. Segev, *et al.*, Dynamic metabolic exchange governs a marine algal-bacterial interaction. *Elife* **5**, e17473 (2016).

16. H. Wang, J. Tomasch, M. Jarek, I. Wagner-Döbler, A dual-species co-cultivation system to study the interactions between *Roseobacters* and dinoflagellates. *Front. Microbiol.* **5**, 311 (2014).
17. H. Wang, *et al.*, Identification of Genetic Modules Mediating the Jekyll and Hyde Interaction of *Dinoroseobacter shibae* with the Dinoflagellate *Prorocentrum minimum*. *Front. Microbiol.* **6**, 1–8 (2015).
18. S. A. Amin, *et al.*, Photolysis of iron-siderophore chelates promotes bacterial-algal mutualism. *Proc. Natl. Acad. Sci.* **106**, 17071–17076 (2009).
19. E. C. Sonnenschein, D. A. Syit, H.-P. Grossart, M. S. Ullrich, Chemotaxis of *Marinobacter adhaerens* and its impact on attachment to the diatom *Thalassiosira weissflogii*. *Appl. Environ. Microbiol.* **78**, 6900–7 (2012).
20. D. H. Green, V. Echavarri-bravo, D. Brennan, M. C. Hart, Bacterial diversity associated with the coccolithophorid algae *Emiliania huxleyi* and *Coccolithus pelagicus f. braarudii*. *Biomed Res. Int.* **2015** (2015).
21. E. L. Harvey, *et al.*, A Bacterial Quorum-Sensing Precursor Induces Mortality in the Marine Coccolithophore, *Emiliania huxleyi*. *Front. Microbiol.* **7**, 59 (2016).
22. L. S. Collier, N. N. Nichols, E. L. Neidle, benK encodes a hydrophobic permease-like protein involved in benzoate degradation by Acinetobacter sp. strain ADP1. *J. Bacteriol.* **179**, 5943–5946 (1997).
23. Y. Nishikawa, Y. Yasumi, S. Noguchi, H. Sakamoto, J. Nikawa, Functional Analyses of *Pseudomonas putida* Benzoate Transporters Expressed in the Yeast *Saccharomyces cerevisiae*. *Biosci. Biotechnol. Biochem.* **72**, 2034–2038 (2008).
24. L. S. Collier, G. L. Gaines, E. L. Neidle, Regulation of Benzoate Degradation in *Acinetobacter* sp. Strain ADP1 by BenM, a LysR-Type Transcriptional Activator. *J. Bacteriol.* **180**, 2493–2501 (1998).
25. R. F. M. van Gorcom, *et al.*, Isolation and molecular characterisation of the benzoate-para-hydroxylase gene (*bphA*) of *Aspergillus niger*: A member of a new gene family of the cytochrome P450 superfamily. *Mol. Gen. Genet. MGG* **223**, 192–197 (1990).
26. P. C. Brzostowicz, A. B. Reams, T. J. Clark, E. L. Neidle, Transcriptional Cross-Regulation of the Catechol and Protocatechuate Branches of the β-Ketoadipate Pathway Contributes to Carbon Source-Dependent Expression of the *Acinetobacter* sp. Strain ADP1 *pobA* Gene. *Appl. Environ. Microbiol.* **69**, 1598–1606 (2003).
27. M. E.-S. Mohamed, A. Zaar, C. Ebenau-Jehle, G. Fuchs, Reinvestigation of a New Type of Aerobic Benzoate Metabolism in the Proteobacterium *Azoarcus evansii*. *J. Bacteriol.* **183**, 1899–1908 (2001).
28. J. Gescher, A. Zaar, M. Mohamed, H. Schägger, G. Fuchs, Genes Coding for a New Pathway of Aerobic Benzoate Metabolism in *Azoarcus evansii*. *J. Bacteriol.* **184**, 6301–6315 (2002).
